# Rapid *in vivo* multiplexed editing (RIME) of the adult mouse liver

**DOI:** 10.1101/2022.03.04.483011

**Authors:** Takeshi Katsuda, Hector Cure, Kamen P. Simeonov, Zoltan Arany, Markus Grompe, Ben Z. Stanger

**Affiliations:** Department of Medicine, University of Pennsylvania, Philadelphia, PA, USA; Department of Cell and Developmental Biology, University of Pennsylvania, Philadelphia, PA, USA; Abramson Family Cancer Research Institute, University of Pennsylvania, Philadelphia, PA, USA; Department of Biomedical Sciences, School of Veterinary Medicine, University of Pennsylvania, Philadelphia, PA, USA; Cardiovascular Institute, Perelman School of Medicine, University of Pennsylvania, Philadelphia, PA, USA; Department of Pediatrics, Oregon Health & Science University, Portland, OR, USA

## Abstract

**Background & Aims:** Assessing mammalian gene function *in vivo* has traditionally relied on manipulation of the mouse genome in embryonic stem cells or peri-zygotic embryos. These approaches are time consuming and require extensive breeding when simultaneous mutations in multiple genes is desired. The aim of this study is to introduce a <u>R</u>apid *<u>I</u>n vivo* <u>M</u>ultiplexed Editing (RIME), and to provide a proof-of-concept of this system.

**Approach & Results:** RIME, a system wherein CRISPR/Cas9 technology, paired with adeno-associated viruses (AAVs), permits the inactivation of one or more genes in the adult mouse liver. The method is quick, requiring as little as 1 month from conceptualization to knockout (KO), and highly efficient, enabling editing in >95% of target cells. To highlight its utility, we used this system to inactivate, alone or in combination, genes with functions spanning metabolism, mitosis, mitochondrial maintenance, and cell proliferation.

**Conclusion:** RIME enables the rapid, efficient, and inexpensive analysis of multiple genes in the mouse liver *in vivo*.

## Introduction

In the last decade, clustered regularly interspaced short palindromic repeats (CRISPR)/Cas9 technology has evolved as a powerful approach for generating genetically engineered mice, enabling researchers to circumvent the time-intensive processes associated with conventional approaches (e.g. embryonic stem cells). In addition, CRISPR/Cas9 has proven to be an efficient tool for reverse genetic screening both *in vitro* (1,2) and *in vivo* (3,4). To overcome challenges associated with the delivery of the Cas9 endonuclease and gene-specific single guide RNAs (sgRNAs), a Cre recombinase-dependent Rosa26-Cas9 knock-in mouse was developed (5), which has facilitated *in vivo* genetic screening in several mouse tissues (6–8). Here, we describe modified protocols that make feasible the editing of single or multiple genes in the adult mouse liver for rapid (<2 month) assessment of gene activity. As proof of concept for this <u>R</u>apid *In vivo* <u>M</u>ultiplexed <u>E</u>diting (RIME) system, we investigated genes involved in mitosis, maintenance of mitochondrial DNA, and cell proliferation, including several whose function in the adult liver has not been previously assessed.

## Materials and Methods

### Mice

Rosa-LSL-Cas9-EGFP mice (5) on c57bl/6J background were purchased from Jackson laboratory (strain #026175) and maintained as homozygotes. 6-8 week-old mice were retro-orbitally injected with various AAV8-EV, AAV8-U6-sgRNA-TBG-Cre or AAV8-PTG-TBG-Cre viruses at 5 × 10^11^ genome copies/mouse. All mouse experiment procedures used in this study were performed following the NIH guidelines. All mouse procedure protocols utilized in this study were in accordance with, and with the approval of, the Institutional Animal Care and Use Committee of the University of Pennsylvania.

### Plasmids and cloning

AAV8-Rep/Cap and Ad5-Helper plasmids were provided from Grompe lab. The backbone vector (empty vector, EV) AAV-U6-sgRNA-TBG-Cre was cloned by replacing the pCBh promoter in the AAV:ITR-U6-sgRNA(backbone)-pCBh-Cre-WPRE-hGHpA-ITR (Addgene #60229). For replacement of the promoter, TBG promoter was PCR-cloned from the pAAV.TBG.PI.eGFP.WPRE.bGH (Addgene #105535) using primers with XbaI cloning site (5’-GGTTCTAGATGCATGTATAATTTCTACAG) and AgeI cloning site (5’-GTCCATGCACTTGTCGAGGTC). For sgRNA cloning, sense and anti-sense oligo DNAs (IDT) (**Table S1**) were phosphorylation using T4 Polynucleotide Kinase (NEB) at 37 °C for 30 minutes, annealed by ramp-down reaction from 95 °C to 25 °C at 5 °C/min, and cloned into the backbone by Golden Gate Assembly using SapI restriction enzyme (NEB) and T4 DNA ligase (NEB). Transformation was performed using into Stbl3 bacteria (Thermo). After Saner-sequencing (Penn DNA Sequencing Core), validated clones were amplified in a large scale (150 ml) and the plasmid DNA was isolated using ZymoPURE Plasmid Maxiprep kit (Zymo Research). Endotoxin was eliminated by treating the plasmid with Endozero columns (Zymo Research).

For the PTG system, we constructed an AAV-hU6-sgEGFP-tRNA-Hnf4a-sg3-TBG-Cre (PTG-*EGFP*/*Hnf4a* in short) and AAV-hU6-*Lats1*-sg1-tRNA-*Lats2*-sg2-tRNA-*Lats2*-sg1-tRNA-*Lats2*-sg2-TBG-Cre (PTG-*Lats1*/*2* in short) by Gibson Assembly. For PTG-*EGFP*/*Hnf4a* cloning, a DNA block containing sg*EGFP*-tRNA-*Hnf4a*-sg2 was PCR-amplified using pGTR plasmid (Addgene #63143) as a template and primers listed in **Table S2** (see also **Fig. S4C**). For PTG-*Lats1*/*2* cloning, 3 DNA blocks (*Lats1*-sg1-tRNA-*Lats1*-sg2, *Lat1*-sg2-tRNA-*Lats2*-sg1, *Lats2*-sg1-tRNA-*Lats2*-sg2) were PCR-amplified using pGTR plasmid and primers listed in **Table S2** (see also **Fig. S5B**). Then, DNA block(s) were assembled into the SapI-linearized AAV vector using the NEBuilder (NEB).

### AAV preparation

90-100% confluent 293T cells in 15 cm dishes were replenished with 15 ml fresh DMEM (Thermo) supplemented with 2% FBS (EMSCO/FISHER) without antibiotics. For a 15 cm plate, 16 μg AAV8-Rep/Cap plasmid, 16 μg Ad5-Helper plasmid, 16 μg AAV-U6-sgRNA-TBG-Cre plasmid and 144 μl of 1 mg/ml Polyetherimide (PEI) (Polysciences) was mixed in 9 ml OptiMEM (Thermo). To target a gene, we typically co-transfected 3 sgRNAs to maximize the KO efficiency. For this end, we combined 5.3 μg of each sgRNA plasmid so that 16 μg in total was co-transfected to 293T cells. To generate AAV-*EGFP*-sg1, EV, AAV-*Hnf4a*-sg2, and PTG viruses, we used 16 μg of single transfer vector. After incubation at room temperature (RT) for 15 minutes, plasmid/PEI complex was added to 293T cells in a dropwise manner, and the plates were gently shaken back and forth to mix the medium evenly. After incubation in a CO2 incubator for 6 days, the cells and culture supernatant were harvested into 50 ml tubes, and centrifuged at 1900 ×*g* for 15 min. The supernatant was transferred to new tubes, and 1/40,000 volume of Benzonase (Sigma-Aldrich) was added and mixed thoroughly by inversion. After digestion of non-viral DNA by incubating at 37 °C for 30 minutes, virus medium was centrifuged at 1900 ×*g* for 15 min, and the supernatant was filtered with 0.22 μm a filter unit with PES membrane (Thermo). Then, 1/4 volume of 40% polyethylene glycol 8000 (PEG8000) in 2.5 M NaCl was added and mixed thoroughly by inversion. Following incubation at 4 °C for at least overnight, precipitated AAV was collected by centrifugation at 3000 ×*g* for 15 min. After removal of the supernatant, precipitate was homogenized in 100 μl PBS per 15 cm dish by through pipetting. Non-AAV precipitate was eliminated by centrifugation at 2200 ×*g* for 5 minutes. Smaller debris were further removed by filtrating the eluted AAV with a 0.45 μm filter columns. This crude AAV was titrated by qPCR using the AAV8-TBG-Cre (Penn Vector Core) as a standard and primers listed in **Table S3**, and directly used for KO experiments without further purification.

### Western blotting

Liver tissue was mechanically homogenized on ice with a pellet pestle (Fisher) prior to lysis. Isolated hepatocytes or homogenized liver tissue were suspended at approximately 1 to 20 volume ratio in RIPA buffer supplemented with Halt™ Protease and Phosphatase Inhibitor Cocktail (Thermo), and pipetted thoroughly. After incubation on ice for 30-60 minutes, lysate was centrifuged at 14,000 ×*g* at 4 °C for 15 minutes, and the supernatant was transferred to new tubes. Protein was then quantified using Pierce™ BCA Protein Assay Kit (Thermo). Equal amounts of protein, 5-20 μg dependent on target abundance, were loaded and separated in 4–20% Mini-PROTEAN gels (Bio-rad), and then transferred to PVDF membrane (Millipore) at 100 V for 90 minutes. After blocking in 5% milk in PBS-T at RT for 30 minutes, membrane was probed with primary antibodies (**Table S4**) at RT for 1 hour or 4 °C overnight. Secondary antibody reaction was performed with donkey anti-rabbit, mouse or goat antibodies conjugated with HRP (Jackson Immuno) at RT for 1 hour. Blots were developed with ECL (Thermo), ECL Plus (Thermo) or West Femto SuperSignal (Thermo) substrate, and imaged using a ChemiDoc Imaging System (Bio-rad).

### Immunofluorescence

Tissue samples were fixed with Zinc Formalin Fixative, pH 6.25 (Polysciences) and embedded in paraffin. Following dewaxing and rehydration, heat-induced epitope retrieval was performed by boiling the specimens in R-buffer (Electron Microscopy Sciences) at 121 °C for 15 min. Then, the specimens were permeabilized with 0.1% Triton X-100 (Fisher). After treatment with the Blocking One Histo at RT for 10 min, the specimens were incubated with primary antibodies (**Table S5**) diluted in 1/20× Blocking One Histo at RT for 1 hour or at 4 °C overnight. The sections were then stained using donkey anti-rabbit, rat or goat antibodies conjugated with AlexaFluor488 or AlexaFluor 594 (Thermo) at 1/300 dilution as well as DAPI (Thermo) at 1/1000 dilution. After incubation at RT for 1 hour, the specimens were mounted in Aqua-Poly/Mount (Polysciences), and imaged on an Olympus IX71 inverted fluorescent microscope.

### F-Actin staining

F-Actin was stained by incubating specimens with Flash Phalloidin™ Red 594 (BioLegend) at 1/500 dilution.

### HE and Oil Red O staining

HE staining was performed by Penn Molecular Pathology and Imaging Core (MPIC), or in-house. When performed in-house, following dewaxing and rehydration, specimens were stained with Harris Hematoxylin (Sigma) for 6-8 minutes. After washed in tap water, excess hematoxylin was removed by dipping in 0.5% HCl/70% EtOH several times. Then, slides were washed in running tap water for 15 minutes, and stained in Eosin (Leica Biosystems Richmond) for 10-20 seconds. After dehydration and EtOH and xylene, specimens were mounted in Permount™ Mounting Medium (Fisher). Oil Red O staining was performed by Penn MPIC using frozen sections.

### Hepatocyte isolation

Livers were perfused with 40 ml of HBSS (Thermo), followed by 40 ml HBSS with 1 mM EGTA (Sigma), and 40 ml of HBSS with 5 mM CaCl_2_ (Sigma) and 40 μg/ml liberase (Sigma). Following perfusion, livers were mechanically dispersed with tweezers, resuspended in 10 ml wash medium (DMEM supplemented with 5% FBS), and filtrated with a 70 μm cell strainer. The cells were centrifuged at 50 ×*g* at 4 °C for 5 minutes. Then, the cells were resuspended in complete percoll solution (10.8 ml percoll (Cytiva), 12.5 ml wash medium and 1.2 ml 10× HBSS per mouse), and centrifuged at 50 ×*g* at 4 °C for 10 minutes. After washed in 10 ml wash medium once by centrifugation at 50 ×*g* at 4 °C for 5 minutes, the cells were used for downstream experiments.

### GFP flow cytometry

Cells were resuspended in flow buffer, HBSS with pH 7.4 and supplemented with 25 mM HEPES (Thermo), 5 mM MgCl_2_ (MedSupply Partners), 1× Pen/Strep (Thermo), 1× Fungizone (Thermo), 1× NEAA (Thermo), 1× Glutamax (Thermo), 0.3% glucose (Sigma), 1× sodium pyruvate (Thermo). After adding 1 μg/ml DAPI (Thermo), GFP fluorescence was analyzed on an LSR II flow cytometer (BD).

### Ploidy analysis

For cellular ploidy analysis, freshly isolated hepatocytes were suspended in 10% FBS-DMEM supplemented with 15 μg/mL Hoechst 33342 (Thermo) and incubated at 37 °C for 30 minutes. Following adding TO-PRO-3 iodide (Thermo) at 1 μM, the cells were analyzed on an LSR II flow cytometer. For nuclear ploidy analysis, nuclei were isolated by suspending cells in nuclei isolation buffer (NIB: pH 7.5, 15 mM Tris-HCl, 60 mM KCl, 15 mM NaCl, 5 mM MgCl_2_, 1 mM CaCl_2_ and 250 mM sucrose) supplemented with 0.1% NP-40 (Millipore) at 10:1 ratio volume on ice for 5 minutes. After 2 rounds of wash by resuspending the nuclei in NP-40-free NIB at 10:1 volume ratio followed by centrifugation at 600 ×*g* for 5 minutes at 4 °C, nuclei were resuspended in 500 μl NIB containing 1 μg/ml DAPI, and analyzed on LSR II.

### Primary culture of hepatocytes

Culture medium was DMEM/F12 (Corning) containing 2.4 g/L NaHCO_3_ and L-glutamine, which was supplemented with 5 mM HEPES (Corning), 30 mg/L L-proline (Sigma), 0.05% BSA (Sigma), 10 ng/ml epidermal growth factor (Sigma), insulin-transferrin-serine (ITS)-X (Thermo), 10^−7^ M dexamethasone (Sigma), 10 mM nicotinamide (Sigma), 1 mM ascorbic acid-2 phosphate (Wako), and gentamycin (Thermo), 10% FBS, 10 μM Y-27632 (LC Laboratories), 0.5 μM A-83-01 (AdooQ BioScience), and 3 μM CHIR99021 (LC Laboratories). Isolated cells were plated to 12-well collagen-coated plates (IWAKI) at 2 × 10^4^ cells/well. Culture medium was replaced every 2-3 days.

### Total RNA isolation and reverse transcription

Total RNA was extracted using NucleoSpin RNA Kit (Takara) following the manufacturer’s instruction. Approximately 500 ng of RNA was reverse transcribed in 20 ul volume using High Capacity cDNA Reverse Transcription Kit (Thermo). cDNA was diluted at 1/20 ratio in water and used for qPCR.

### Total DNA isolation

Total DNA was isolated from hepatocytes using DNeasy Blood & Tissue Kit (Qiagen). DNA was diluted to 5-10 ng/μl and used for qPCR.

### qPCR

qPCR was performed at 10 μl/well using Bio-Rad CFX 384 qPCR machine (Bio-rad): 3 μl cDNA or genomic/mitochondrial DNA diluted in water as described above, 0.25 μl each of 10 μM forward and reverse primers (**Table S3**), 1.5 μl H_2_O and 5 μl SsoAdvanced SYBR reagent (Bio-rad).

### Transmission electron microscopy

Electron microscopy imaging was performed at Penn Electron Microscopy Resource Laboratory.

### Blood chemistry

All the blood chemistry was tested by IDEXX laboratories.

## Results

### Efficient hepatocyte knockout is achieved within 2 weeks after AAV injection

The RIME system has two components: (i) an adeno-associated virus serotype 8 (AAV8) carrying *Cre* recombinase under the control of a tissue specific promoter and a single guide RNA (sgRNA) under the control of the human U6 promoter, and (ii) host LSL-Cas9-EGFP mice (5), in which CAG-driven expression of *Cas9* and *EGFP* is activated by Cre-mediated removal of an upstream transcriptional stop sequence (**Fig. 1A**). To target hepatocytes, we modified the existing AAV construct (5) by replacing the ubiquitous CBh promoter with the hepatocyte-specific thyroid hormone-binding globulin (TBG) promoter to drive Cre expression. AAV preparation from HEK293T cells was performed using a streamlined protocol (see Methods), and crudely processed viral preps were introduced into LSL-Cas9-EGFP mice by retro-orbital injection.

**Figure 1.**
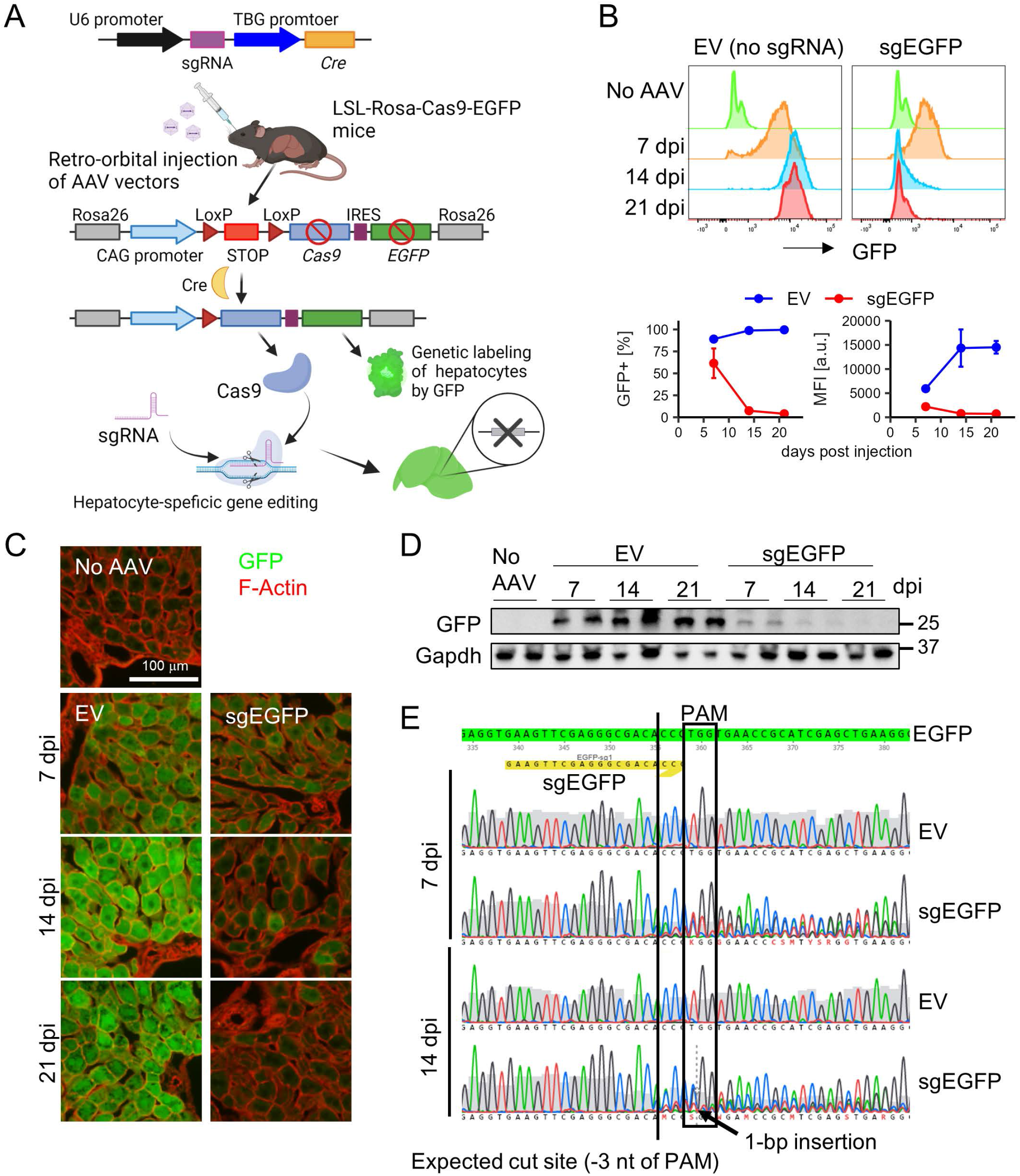
Efficient knockout in hepatocytes by AAV-mediated delivery of sgRNA to Cas9-EGFP mice. (A) Schematic representation of the RIME strategy. (B) Flow cytometric evaluation of knockout efficiency of EGFP as a function of days post injection (dpi) of AAVs (n = 2-3 at each time point). (C) Representative images of co-staining of GFP and F-Actin at designated time points. (D) Western blots of EGFP at the designated time points. (E) Representative chromatograph of Sanger sequenced PCR amplicons capturing the expected cut sites.

We first estimated the kinetics of gene editing by targeting *EGFP*. Injection of AAV8 lacking an sgRNA (empty vector, AAV-EV) resulted in the expression of EGFP in ~90% hepatocytes at 7 days post injection (dpi) and >99% at 14 dpi, as assessed by flow cytometry (**Fig. 1B**). By contrast, injection of AAV8 packaged with an sgRNA targeting EGFP (AAV-sg*EGFP*) led to substantial decrease of GFP+ cell fraction as well as the mean fluorescent intensity of the GFP+ cells at 14-21 dpi (**Fig. 1B**). This observation was further validated by immunohistochemistry (IHC) (**Fig. 1C**) and western blotting (WB) (**Fig. 1D**). Sanger sequencing of bulk genomic DNA derived from isolated hepatocytes confirmed perturbation in the chromatograph at the proximity of the expected sg*EGFP* cut site (**Fig. 1E**). Thus, gene editing in this system is rapid and highly efficient.

### Targeting *Hnf4a* in the adult liver

To assess our system functionally, we designed AAV8 vectors with sgRNAs targeting *Hnf4a*, a known master regulator of hepatocyte identity and function. In a previous study, in which *Alb*-CreER^T2^ mice were crossed to *Hnf4a^loxP/loxP^* mice, *Hnf4a* deletion was noted to result in marked hepatocyte proliferation and hypertrophy within 2-3 weeks of tamoxifen treatment, contrasting with the phenotypes observed following embryonic deletion of *Hnf4a* (9). We therefore reasoned that knocking out *Hnf4a* (*Hnf4a* KO) in adult hepatocytes would provide a good test of the system’s fidelity.

We generated a pooled AAV8 viral prep carrying 3 sgRNAs against *Hnf4a* by co-infecting HEK293T cells with plasmids carrying sgRNAs targeting different parts of the gene (**Fig. S1A**). At 10 dpi, we observed a decrease in *Hnf4a* mRNA *in vivo* by qRT-PCR with primer pairs proximal to each cut site (**Fig. S1B**) and confirmed complete loss of Hnf4a protein by WB (**Fig. 2A**). Accordingly, the liver of the mice injected with sg*Hnf4a* became pale, a characteristic of steatosis (**Fig. 2B**). Hematoxylin and eosin (HE) staining revealed scant cytoplasm, and oil Red O staining confirmed abnormal accumulation of oil droplets in the sg*Hnf4a*-injected hepatocytes (**Fig. 2C**). Blood chemistries indicated that liver damage peaked at 10-14 dpi, with recovery evident by 22 dpi (**Fig. 2D**), observations consistent with the finding that bulk *Hnf4a* mRNA expression began to recover by 3 wpi (**Fig. S1B**) in alignment with previous work (9). Notably, we found that the expression of several biliary epithelial cell (BEC) genes increased in *Hnf4a*-KO livers (**Fig. 2E**) while the expression of several hepatocyte-specific genes decreased (**Fig. S1C**). These results suggest that *Hn4a*-KO hepatocytes engage in hepatocyte-to-biliary reprogramming (10,11) and/or that the liver injury associated with *Hnf4a* loss induced a ductal cell expansion and reduced hepatocyte gene expression.

**Figure 2.**
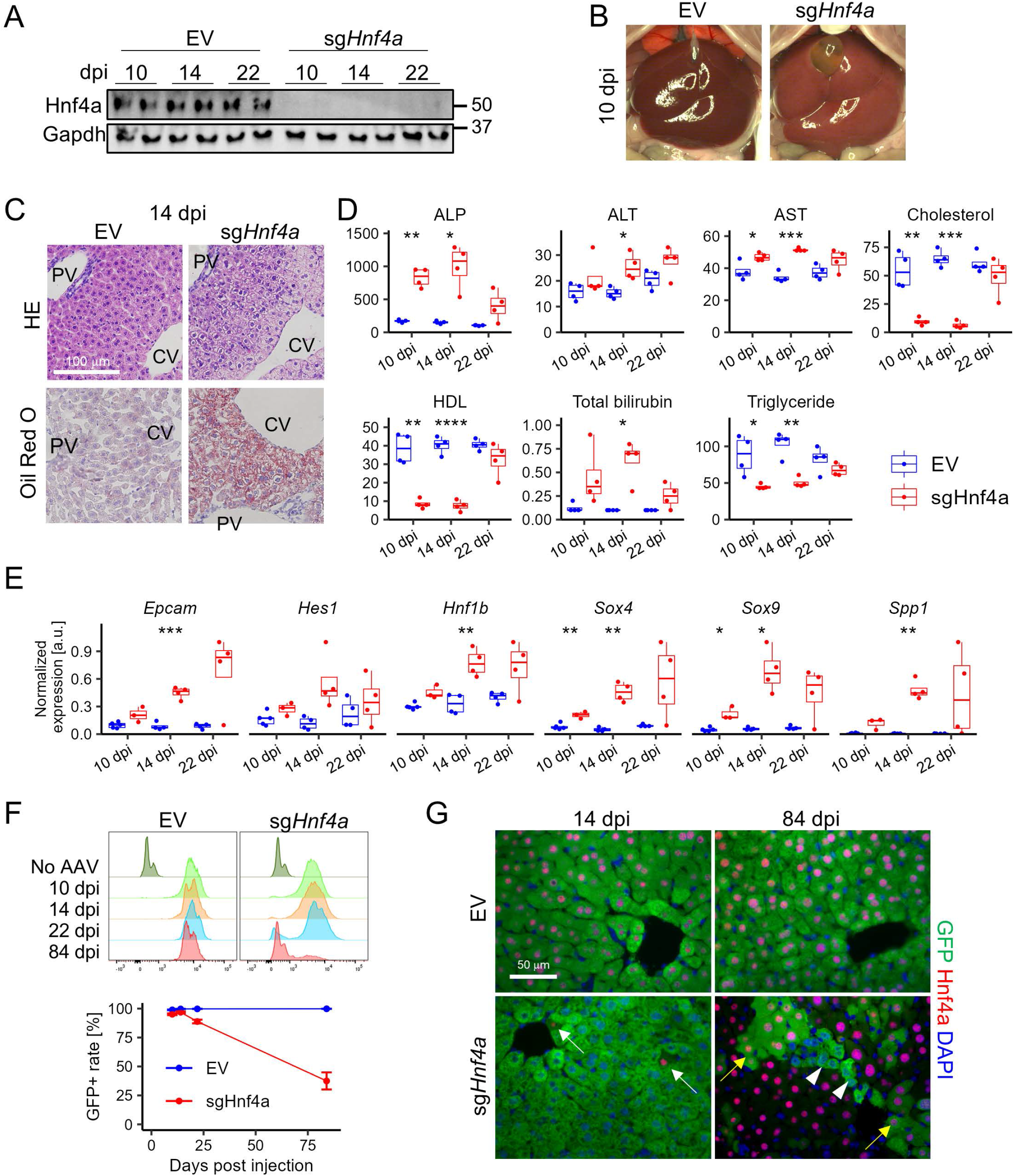
*Hnf4a*-LKO causes liver injury and repopulation by escaper cells. (A) Western blots of Hnf4a at the designated time points. (B) Representative macroscopic images of EV or sg*Hnf4a*-injected mouse livers at 10 dpi. (C) Representative images of HE (top) and Oil Red O staining (bottom) at 14 dpi. PV and CV indicate portal vein and central vein, respectively. (D) Blood biochemistries of EV or sg*Hnf4a*-injected mice (n = 4 at each time point). The units on the y-axes are U/L for ALP, ALT and AST, and mg/dL for Cholesterol, HDL, Total bilirubin and Triglyceride. (E) Gene expression analysis of biliary epithelial cell marker genes by qRT-PCR (n = 4 at each time point). The data are normalized to *Gapdh* expression. (F) Quantification of GFP+ cells by flow cytometry (n = 4-6 at each time point). (G) Representative immunofluorescent images for EV or sg*Hnf4a*-injected livers at the designated time points. White arrows indicate GFP-Hnf4a+ cells. Yellow arrows indicate GFP+Hnf4a+ cells. Arrow heads indicate GFP+Hnf4a-cells. Statistical differences were calculated using multiple t-test. *p < 0.05, **p < 0.01, ***p < 0.001, ****p < 0.0001

To obtain insight into the origin of the cells contributing to recovery of *Hnf4a* expression, we measured the abundance of EGFP-labeled cells by flow cytometry. While the vast majority of hepatocytes were EGFP+ at 14 dpi (96.9 ± 0.71%), the peak of labeling, *Hnf4a* KO mice exhibited a decrease in the EGFP+ fraction as early by 22 dpi (88.9 ± 2.0%) and a further decrease at 84 dpi (37.5 ± 7.4%) (**Fig. 2F**), raising the possibility that hepatocytes that had escaped *Hnf4a* deletion were responsible for repopulating the liver at the expense of *Hnf4a* deficient cells. IHC analysis at 84 dpi confirmed that most of the hepatocytes in *Hnf4a* KO livers expressed Hnf4a, whether they were EGFP+ or EGFP-(**Fig. 2G**). Thus, cells that escaped either Cre-mediated recombination or Cas9-mediated editing of the *Hnf4a* gene were the dominant source of hepatocytes more than 2 months after infection. As these results recapitulated and expanded findings previously obtained through traditional gene targeting approaches (see Discussion), we proceeded to target genes whose functions have not previously been investigated in the liver.

### *Kif11* is necessary for the maintenance of hepatocyte ploidy

Approximately 70-90% of adult hepatocytes are polyploid (12,13). Polyploidization of hepatocytes occurs mainly through cytokinesis failure and is developmentally programmed to begin at weaning and increase with age (13). Genetic studies have identified multiple cell cycle-related genes as suppressors or promoters of polyploidization in hepatocytes, including members of the E2F family (reviewed in (12)). Deletion of E2F family members can lead to either an increased (*E2f1*, *E2f2* and *E2f3*) or a decreased (*E2f7* and *E2f8*) number of polyploid hepatocytes through suppression or promotion of endoreplication, respectively. However, these genes have versatile functions in cell cycle regulation, leaving the detailed molecular mechanism of hepatocyte polyploidization unclear.

To evaluate whether disrupting mitotic spindle formation would affect the ploidy status of quiescent hepatocytes, we targeted the *Kif11* (kinesin family member 11) gene. *Kif11*, also known as kinesin-5, is essential for mitosis, where it participates in the self-assembly of the microtubule-based mitotic spindle (14). Disruption of *Kif11* in dividing cells results in mitotic catastrophe and cell death, and thus germline KO of *Kif11* causes early embryonic lethality (15). *Kif11* also contributes to axon extension in adult neurons (14), but its role in other tissues – particularly in quiescent cells – remains unknown.

We injected AAV-EV or AAV-*sgKif11* viruses into 6-week-old LSL-Cas9-EGFP mice to generate *Kif11* KO mice (or EV controls) and harvested hepatocytes after 2 weeks. Neither EV nor *Kif11* KO hepatocytes had detectable levels of Kif11 protein by western blot (**Fig. 3A**), consistent with quiescent state of most hepatocytes in the normal adult liver (16). However, upon plating in optimized hepatocyte culture medium (17), EV hepatocytes readily expressed Kif11, whereas *Kif11* KO hepatocytes failed to express the protein, confirming efficient deletion (**Fig. 3A**). Nevertheless, *Kif11* KO hepatocytes remained viable and expanded in culture, exhibiting enlarged nuclei (**Fig. 3B**).

**Figure 3.**
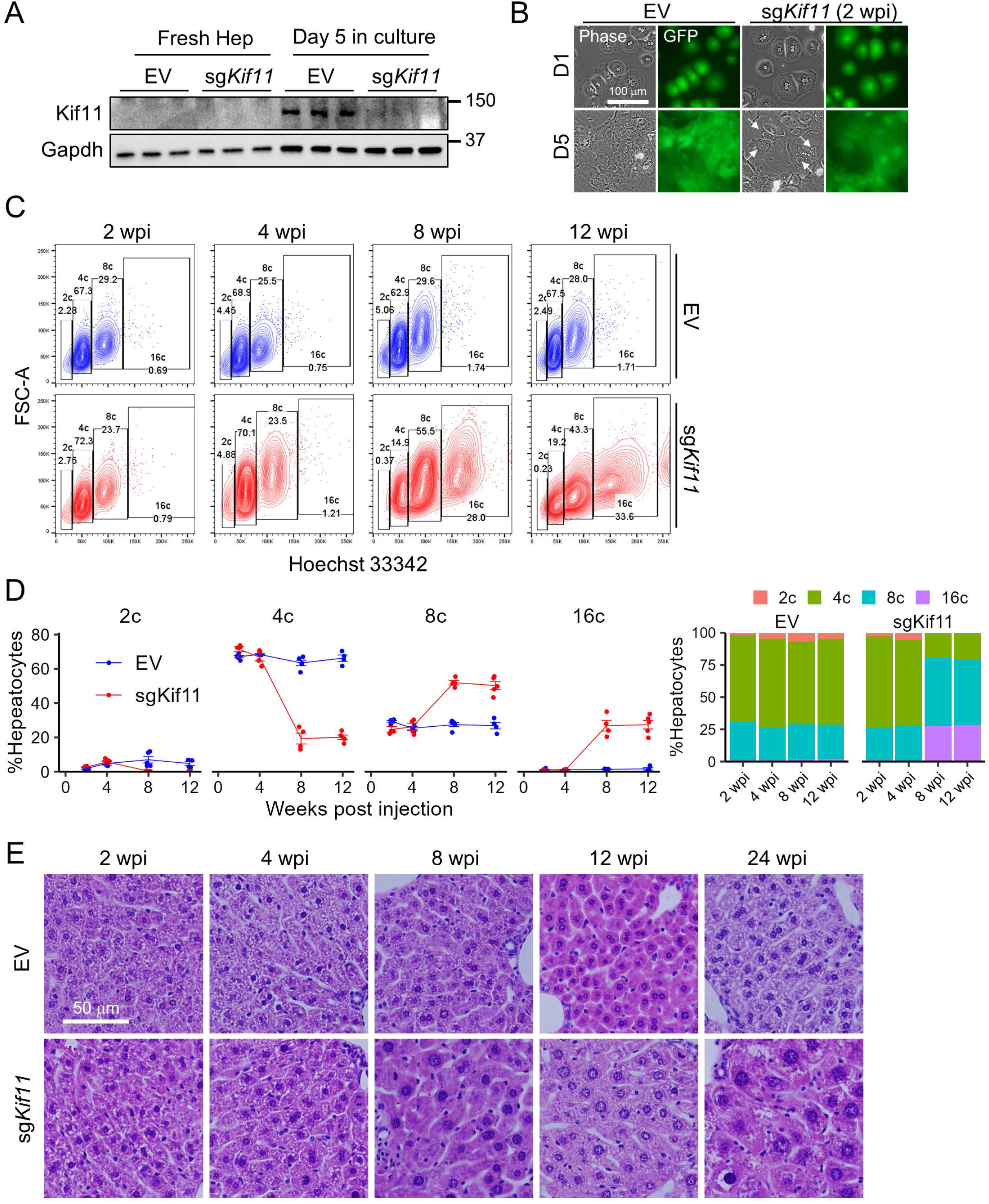
LKO of *Kif11*, a gene involved in mitotic spindle formation, causes elevation of hepatocyte ploidy. (A) Western blots of freshly isolated hepatocytes and hepatocytes cultured in the presence of three signaling inhibitors – Y27632 (ROCK inhibitor), A83-01 (TGFβ inhibitor) and CHIR99021 (GSK3 inhibitor) (18) – for 5 days. Hepatocytes were isolated from Cas9-EGFP mice injected with EV and sg*Kif11* at 2 wpi. (B) Representative microscopic images of EV or *sgKif11*-injected hepatocytes 1 day and 5 days after plating. Arrows indicate cells with abnormally large nuclei. (C) Representative flow cytometry plots of Hoechst33342-loaded hepatocytes isolated from EV- or *sgKif11*–injected mice at the designated time points. (D) Quantification of data in (C) (n = 3-5 for each data point). Stacked bar plots (far right) indicate the distribution of hepatocytes with different ploidy (data represent the mean of n = 3-5). (E) Representative HE images of EV or *sgKif11*-injected mouse livers at the designated time points.

Next, we measured ploidy in EV and *Kif11* KO hepatocytes by performing Hoeschst staining on cells obtained from mice between 2 and 12 wpi. While EV hepatocytes showed minimal changes in their ploidy profile over the 12 week period, *Kif11* KO hepatocytes exhibited a dramatic increase in 8c and 16c hepatocytes over the same time period, particularly between 4 and 8 wpi (**Figs. 3C,D**). HE staining confirmed that both nuclear size and cell size were increased in hepatocytes from *Kif11* KO livers (**Fig. 3E**). Intriguingly, most hepatocytes in *Kif11* KO livers were mono-nucleated (**Fig. 3E**), suggesting that the increase in ploidy was the result of endoreplication rather than failed cytokinesis (following successful karyokinesis). Flow cytometry and quantification of ploidy in isolated nuclei support this conclusion (**Figs. S2A,B**).

Despite this hepatocyte enlargement, the overall size of the liver in *Kif11* KO mice was unaltered at 13 wpi (**Fig. S2C**), suggesting that total hepatocyte number was reduced. Cell death followed by hepatocyte hypertrophy may be one mechanism, as Miyaoka et al. reported that hepatocytes compensate for mild tissue loss (30% partial hepatectomy) by increasing hepatocyte volume rather than through cell division (18). To determine whether cells that had escaped *Kif11* loss were more fit than *Kif11* KO cells, we again measured the fraction of EGFP+ over time. While most hepatocytes remained EGFP+ through weeks 8 and 12 wpi (EGFP+ rate = 96.6 ± 2.2% and 95.9 ± 1.6% respectively), we observed substantial repopulation by escaper (EGFP-) cells at 24 wpi (EGFP+ rate = 60.4 ± 16.9%) (**Fig. S2D**). These results suggest that loss of Kif11 is detrimental to hepatocyte viability, even under homeostatic conditions where rates of hepatocyte proliferation are low.

### Mice tolerate transient loss of mitochondrial DNA in hepatocytes

Mitochondrial diseases can present at any age and manifest in any organ (19), and germline mutations that disrupt mitochondrial function are the most common cause of mitochondrial diseases (20). Although mitochondria have their own ~16.5kb genome, encoding 37 genes, the genes responsible for maintaining mitochondrial DNA (mtDNA) are encoded in the nucleus. Mutations in mitochondrial maintenance genes result in mtDNA depletion syndrome (MDS), but the mechanisms of mtDNA loss, and the tissue specificity of the resulting disease phenotypes, remain unclear (20). To date, there has been limited investigation of the major mitochondrial maintenance genes in the liver, where disease phenotypes associated with MDS are common.

To address this knowledge gap, we used RIME to identify liver phenotypes associated with loss of three mtDNA maintenance genes – *Tfam*, *Polg* and *Polg2* – germline deletion of which cause embryonic lethality at E8.0-10.5 in the mouse (21–23). *Tfam* was originally identified as a transcription factor necessary for mtDNA transcription and was subsequently found to be important for mtDNA replication and the maintenance *in vitro* (21,24). Heterotrimeric polymerase γ, the mitochondria-specific polymerase, is composed of a catalytic subunit encoded by *Polg* and a homodimeric accessory subunit encoded by *Polg2*, which is required for the efficient binding of Polg to the mtDNA. Defects in all three genes are associated with MDS, but expressivity and tissue tropism vary; specifically, defects in *TFAM* and *POLG* typically involve liver diseases, while liver involvement with *POLG2* mutations is rare (19,20,25).

To inactivate these genes in the liver, we injected LSL-Cas9-EGFP mice with AAV viruses carrying targeting sgRNAs as described above. We first performed western blots to confirm protein loss. While Polg protein was absent by 2 wpi, Tfam persisted longer, possibly due to greater protein or mRNA stability, but was largely absent by 6-8 wpi (**Fig. 4A**). We were unable to detect Polg2 by western blotting using 2 commercially available antibodies (data not shown). As an alternative, we isolated genomic DNA from the AAV-sg*Polg*2-infected hepatocytes and bulk sequenced the region containing the expected cut sites (**Fig. S3A**). This demonstrated chromatographic perturbation specifically in the hepatocytes injected with AAV-sg*Polg2* at each of the 3 cut sites (**Fig. S3A**). In addition, qRT-PCR using the *Polg2*-sgRNAs as a forward primer for each of the 3 cut sites confirmed the reduction of *Polg2* transcripts near the cut sites (**Fig. 4B**). Based on these results, we conclude that all the three genes are susceptible to *in vivo* knockout using this technique.

**Figure 4.**
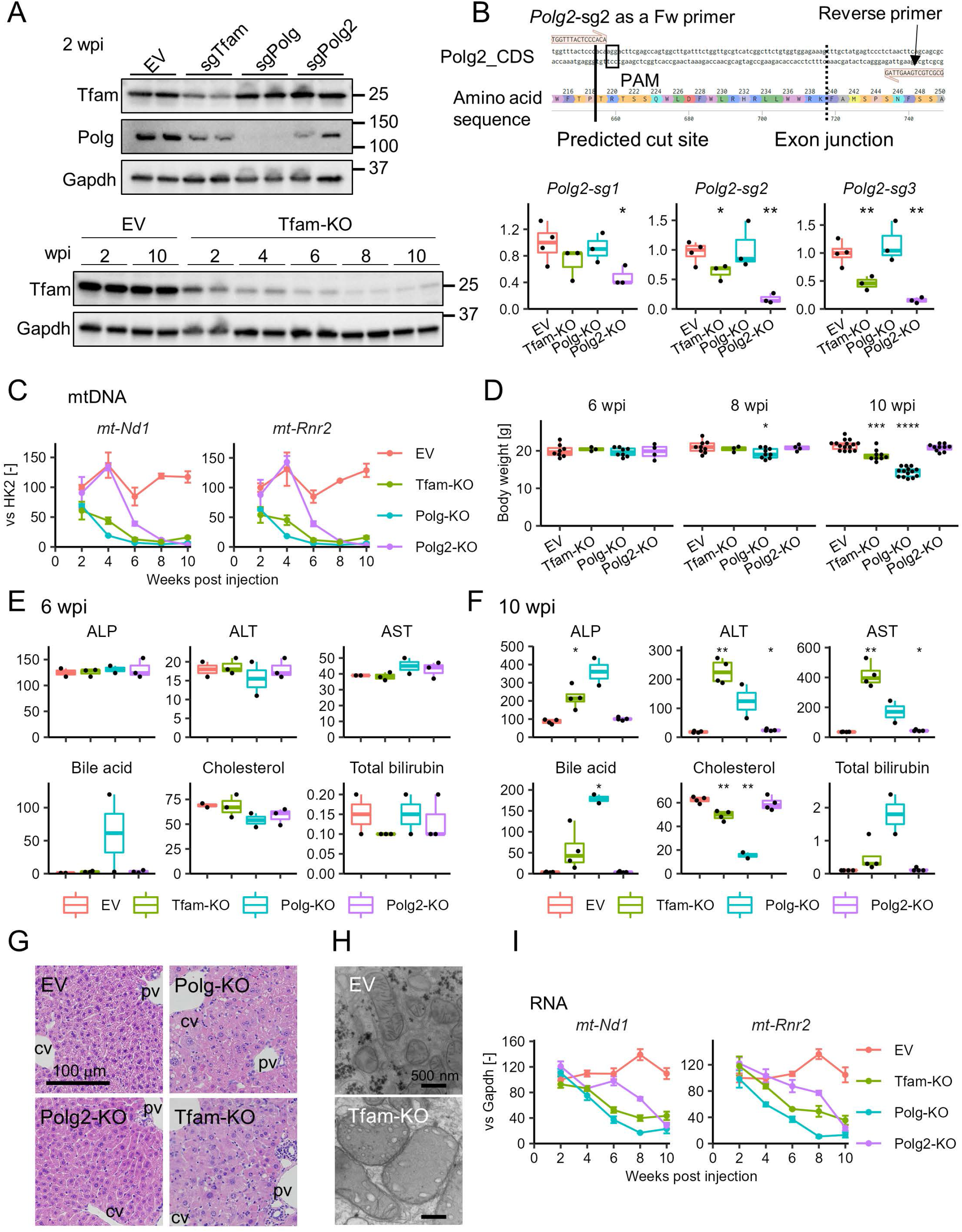
Mice tolerate severe mtDNA loss. (A) KO validation for Tfam and Polg by Western blotting. (B) KO validation for *Polg2* by qRT-PCR using sgRNA as a forward primer. The upper diagram indicates the amplified region targeting the *Polg2*-sg2 cut site as an example. The lower panels indicate the quantification of qRT-PCR targeting at each cut site. (C) Changes in mtDNA abundance in hepatocytes over time as determined by qPCR (n = 2-4 at each point). mtDNA contents are normalized to the *Hk2* gene (nuclear encoded). The values of EV-injected hepatocytes at 2 wpi are set to 100%. (D) Body weights of mice injected with EV, sg*Tfam*, sg*Polg* and sg*Polg2* measured at the designated time points. (E) Blood biochemistry at 6 wpi (n = 2-3). The units on the y-axes are U/L for ALP, ALT and AST, μM for bile acid, and mg/dL for Cholesterol and Total bilirubin. (F) Blood biochemistry at 10 wpi (n = 2-4). The units on the y-axes are U/L for ALP, ALT and AST, μM for bile acid, and mg/dL for Cholesterol and Total bilirubin. (G) HE staining of EV, sg*Tfam*, sg*Polg* and sg*Polg2*-injected liver at 10 wpi. (H) Transmission electron microscopic images of mitochondria of EV and sg*Tfam*-injected hepatocytes at 10 wpi. (I) Time course change in the transcripts of mtDNA-encoded genes in hepatocytes as determined by qPCR (n = 2-4 at each point). Expression levels are normalized to *Gapdh*. The values of EV-injected hepatocytes at 2 wpi are set to 100%. Statistical differences were calculated using multiple t-test with EV as the reference. *p < 0.05, **p < 0.01, ***p < 0.001, ****p < 0.0001.

Unexpectedly, we observed that liver-specific deletion of either *Tfam* or *Polg2* led to a reduction of Polg protein as early as 2 wpi, while knocking out *Polg* or *Polg2* led to mild reductions of Tfam protein (**Fig. S3B**). qRT-PCR did not detect downregulation of *Polg* mRNA by *Tfam*-KO or *Polg2*-KO (Fig. S3C). These results suggest that Tfam and Polg2 are required for Polg stability at the protein level, consistent with a recent report that *Polg2* KO or knockdown *in vitro* leads to reductions in Polg and Tfam (26). We also observed that *Tfam*-KO hepatocytes downregulated *Polg2* mRNA expression (**Fig. S3C**), suggestive of additional regulation at the RNA level. Collectively, these data underscore a highly interdependent regulatory system among these 3 genes.

As *Tfam, Polg* and *Polg2* are involved in mtDNA maintenance, we performed qPCR to measure the abundance (DNA) of several mtDNA-encoded genes, including *mt-Nd1* (NADH-ubiquinone oxidoreductase chain 1) and *mt-Rnr2* (16S ribosomal RNA), in KO livers. Analysis of hepatocytes isolated from *Tfam*-KO and *Polg*-KO mice between 2 wpi and 10 wpi revealed steep declines in mtDNA levels (**Fig. 4C**). Specifically, mtDNA levels decreased by ~60-80% at 4 wpi and by more than 95% at 8 wpi (**Fig. 4C**). Despite this dramatic loss of mtDNA, there were few gross manifestations until 8 wpi, when some animals began to exhibit weight loss (**Fig. 4D**). Likewise, there was little evidence of laboratory abnormalities at 6 wpi, when KO animals had lost anywhere from 60-90% of mtDNA, aside from elevated bile acids in one *Polg* KO animal (**Fig. 4E**). By 10 wpi, however, *Polg* KO and *Tfam* KO animals exhibited signs of hepatocellular death and cholestasis (**Fig. 4F**). Liver injury was confirmed by HE staining (**Fig. 4G**), and transmission electron microscopy of 10 wpi *Tfam* KO livers revealed giant mitochondria with poorly developed cristae (**Fig. 4H**). As expected, GFP-negative “escaper” cells repopulated the livers of *Polg* KO and *Tfam* KO animals, although this did not occur to a significant degree until 13 wpi, well after the manifestation of these phenotypes (**Fig. S3D**). Surprisingly, *Polg2* KO animals exhibited only slight/undetectable changes in body weight (**Fig. 4D**), liver chemistries (**Fig. 4F**) or histology (**Fig. 4G**) at 10 wpi despite the nearly complete loss of mtDNA at that timepoint (**Fig. 4C**). Additional follow-up revealed evidence of liver damage in *Polg2* KO mice a week later, at 11 wpi (**Fig. S3E**).

Given that reductions in Polg levels were was readily and robustly observed in *Tfam* KO and *Polg2* KO hepatocytes (**Fig. S3B**), we sought to determine whether Polg loss is the common cause of the liver injury in all liver knockout (LKO) animals. To this end, we exploited our system’s ability to target multiple genes simultaneously though the co-injection of multiple AAVs. Using this approach, we generated *Polg*/*Polg2* double KO (DKO) and *Tfam*/*Polg*/*Polg2* triple KO (TKO) mice. *Polg*/*Polg2* DKO slightly exacerbated the weight loss at 10 wpi, while the *Tfam*/*Polg*/*Polg2* TKO mice (3 of 3) died at 9-10 wpi (**Fig. S3F**). These results indicate an additive effect of simultaneous KO of these three genes, suggesting that both distinctive and redundant mechanisms underlie the liver injuries caused by their loss.

Given that manifestations of liver damage became apparent only 2 weeks after mtDNA levels dropped below 10%, we hypothesized that persistence of mitochondrially-encoded RNAs might have allowed cells to survive in the face of mtDNA depletion. To test this, we performed a time course of *mt-Rnr2* and *mt-Nd1* RNA expression in the three mutants. As shown in **Fig. 4I**, levels of *mt-Rnr2* and *mt-Nd1* RNA were maintained at 50% of control levels at 6 wpi in *Polg* KO and *Tfam* KO mice and at 8 wpi in *Polg2* KO mice, consistent with the delayed kinetics of liver damage in these mice. These results suggest that hepatocyte function and viability are temporarily preserved in cells lacking mtDNA through the persistence of mitochondrially-encoded RNAs. Prior proteomic studies have estimated that the median half-life of mitochondrial proteins in the liver is 4-5 days (27). Hence, levels of mitochondrially-encoded proteins would be expected to undergo steep reductions between 8-10 wpi (when mRNA levels nadir), a timing that is consistent with the expansion of escaper (EGFP-negative) hepatocytes at 10-13 wpi (**Fig. S3D**). These findings provide insight into the concept of “mitochondrial threshold effects,” wherein failure of the mitochondrial electron transport chain occurs once the frequency of mtDNA mutations exceeds a certain threshold, thought to be 60-80% (28). Our data indicate that hepatocytes can tolerate a more severe and complete loss of mtDNA for several weeks, raising the possibility that the liver has a higher threshold for mtDNA loss compared to other tissues.

### PTG: a multiplexed polycistronic sgRNA system

The ability to target multiple genes simultaneously (**Fig. S3F**) could vastly simplify the time-consuming breeding necessary to generate KO mice carrying multiple mutant alleles. To confirm that such an approach could work, we co-injected AAVs carrying sgRNAs targeting *EGFP* and *Hnf4a* (**Fig. S4A**, left). As predicted, this resulted in livers that were both steatotic and EGFP-negative (**Fig. S4B**). Next, we asked whether it would be possible to modify the system to incorporate multiple sgRNAs into a single viral vector. To do so, we introduced a polycistronic sgRNA expression platform (PTG: <u>p</u>olycistronic tRNA-s<u>g</u>RNA) that utilizes the cell’s tRNA processing machinery (**Fig. S4A**, right**, Fig. S4C**) (see Methods for details). Mice injected with AAV carrying the polycistronic PTG-*EGFP*/*Hnf4a* construct recapitulated features of AAV co-injection, including steatosis and loss of EGFP expression (**Fig. S4B**). Reductions in Hnf4α levels following AAV-PTG-*EGFP*/*Hnf4a* injection were confirmed by western blotting (**Fig. S4D**). Reductions in EGFP levels were confirmed by flow cytometry, although loss of signal (mean fluorescence intensity, MFI) appeared to be slower with the polycistronic vector as compared to co-injection (**Fig. S4D,E**). These findings suggest that multiplexed gene editing *in vivo* can be streamlined with polycistronic viral vectors.

We next sought to apply this system to another biological context. To this end, we chose to knock out the large tumor suppressor homologs 1 and 2 (*Lats1/2*), which regulate liver development by antagonizing the transcriptional coactivators YAP and TAZ (29). We first confirmed that co-injection of AAV-*Lats1*-sg1/2 and AAV-*Lats2*-sg1/2, but not individual injection of *Lats1*-sg1/2 or *Lats2*-sg1/2, resulted in liver overgrowth at 3 wpi (**Fig. 5A-C, Fig. S5A**). In contrast to one prior study, in which animals survived for 2 months after injection of Adenovirus-*Cre* into *Lats1^−/−^;Lats2^fl/fl^* mice (29), no animals survived beyond 4 wpi in our AAV-co-injected *Lats1*/*Lats2* DKO cohort (n = 6).

**Figure 5.**
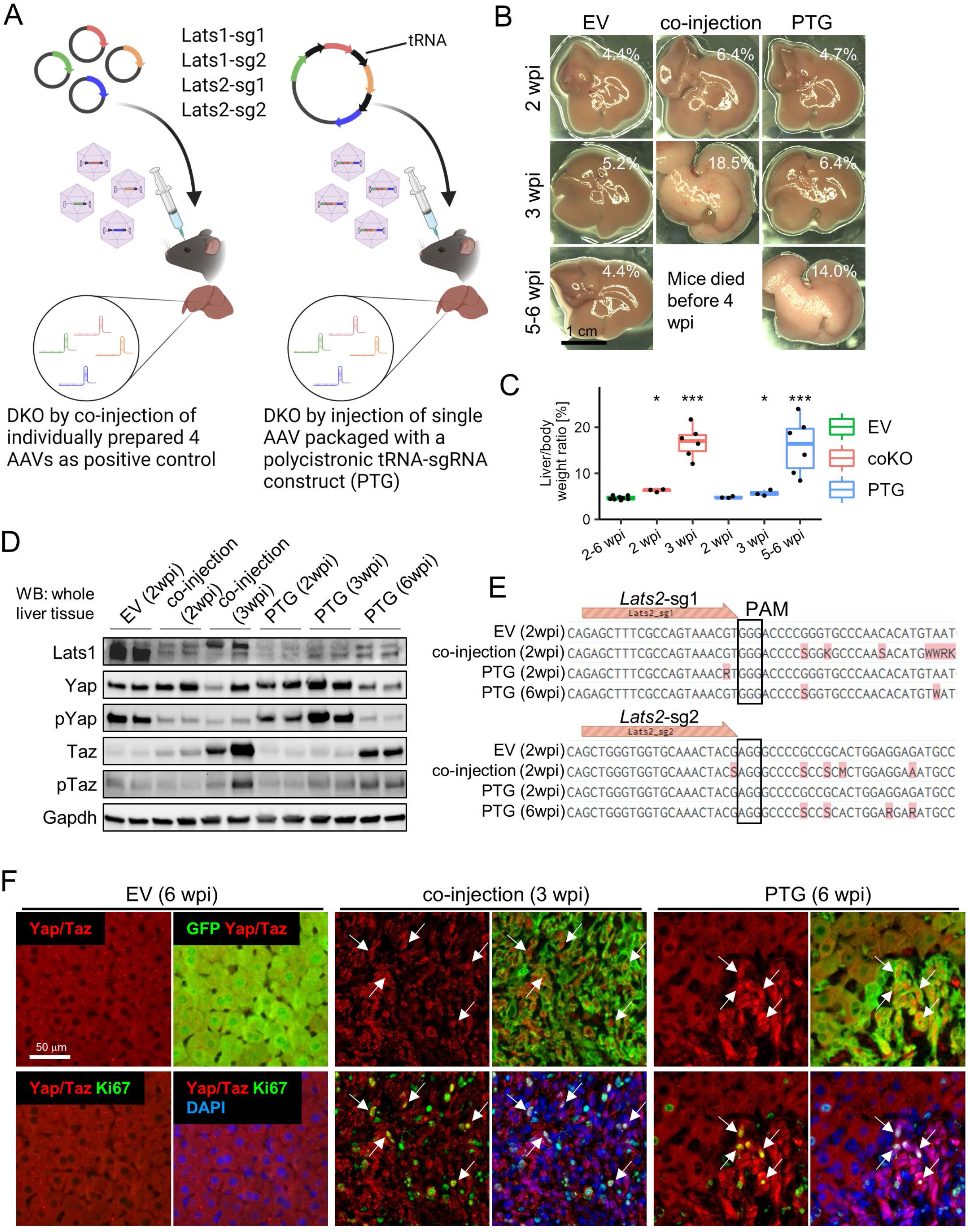
Multiplex mosaic knockout of Hippo-Yap pathway components induce cell competition. (A) Schematic representation of the strategy of DKO of *Lats1*/*2* by co-injection of 4 AAVs (left) and by injection of single AAV-PTG-*Lats1*/*2* (right). (B) Representative macroscopic images of *Lats1*/*Lats2* DKO at the designated time points. Percentages shown in the photos indicate liver/body weight ratios. (C) Liver/body weight ratios at the designated time points are shown for *Lats1*/*Lats2* DKO mice generated by either co-injection or PTG strategies. Mice harvested 2-6 weeks after injection of EV are shown as control. (D) Western blots for validation of KO of *Lats1* and *Lats2* as well as the subsequent influence on Yap and Taz phosphorylation. (E) Bulk Sanger sequencing result for each of the cut sites targeted by 2 *Lats2* sgRNAs. Chromatograph images corresponding to these data are shown in Fig. S6B. (F) Immunofluorescence images for Yap/Taz, GFP and Ki67 using the livers harvested from mice injected with EV, *Lats1*-sg1/2 and *Lats2*-sg1/2 as co-injection and PTG-*Lats1*/*2* at the designated time points. Nuclei were counterstained with DAPI. Arrows indicate the cells co-stained with of nuclei-localized Yap/Taz and Ki67.

Next, we designed a PTG-*Lats1*/*2* vector which incorporated the two *Lats1* sgRNAs and two *Lats2* sgRNAs used in the co-injection experiment (**Fig. 5A**, right, **Fig. S5B**). In contrast to the co-injection cohort, the PTG-*Lats1/2* cohort exhibited only modest liver overgrowth at 3 wpi (**Fig. 5B,C**). By 5-6 wpi, however, liver overgrowth in the PTG-*Lats1/2* cohort was comparable to that of the co-injection cohort at 3 wpi (**Fig. 5B,C**). Western blotting confirmed substantial reductions in Lats1 levels (**Fig. 5D**), but as we were unable to reliably monitor Lats2 levels by western blotting (data not shown), we performed bulk Sanger sequencing to confirm mutations in *Lats2* in co-injected mice at 2 wpi and in PTG mice at 6 wpi (**Fig. 5E, Fig. S5C**). Accordingly, levels of phospho-Yap (pYap) were diminished in the co-injected cohort at 2 wpi and in the PTG cohort at 6 wpi, while total YAP levels did not change significantly (**Fig. 5D**). Unexpectedly, we observed a robust increase in the level of total Taz, but not total Yap, at 3 wpi in the co-injected mice and 6 wpi in PTG mice (**Fig. 5D**), suggesting distinct mechanisms regulating Yap and Taz stabilization by Lats1 and Lats2.

Finally, we compared the histological consequences of *Lats1/2* inactivation in the liver using these two approaches. HE staining as well as immunofluorescence staining with biliary markers revealed the emergence of hepatocyte-derived BEC-like cells in both the co-injection and PTG cohorts (**Fig. S6**), as expected (30). In the control (EV) liver, Yap and Taz staining was limited to the hepatocyte cytoplasm, and hepatocyte proliferation (measured by Ki67) was low (**Fig. 5F**, left). In mice that received a co-injection of AAV-*Lats1*-sg1/2 and AAV-*Lats2*-sg1/2, by contrast, Yap and Taz were detectable in both the nucleus and cytoplasm, and hepatocyte proliferation was elevated (**Fig. 5F**, middle). Similar effects were observed in the PTG-*Lats1/2* cohort, although more normal regions were observed as compared to the co-injection cohort (**Fig. 5F**, right). In summary, the PTG strategy permits mosaic inactivation of multiple genes in the mouse liver.

## Discussion

This report describes RIME, a suite of tools that facilitate the inactivation of single or multiple genes in the adult mouse liver. To demonstrate the feasibility and utility of this system, we knocked out seven genes with functions spanning metabolism, mitosis, mitochondrial maintenance, and cell proliferation. As our approach is streamlined for viral production and utilizes an LSL-Cas9-EGFP mouse colony, most of these experiments were able to move from concept to realization within a month, with evidence of gene inactivation occurring 1-2 weeks after injection of AAVs. These features allowed us to efficiently evaluate the function of the *Kif11* gene, whose role in hepatocyte ploidy has not been previously studied (**Fig. 3, Fig. S2**).

In addition to its speed, RIME provides other advantages. For example, the system contains a built-in lineage trace (the result of Cre-mediated expression of an EGFP reporter), a feature which allows a re-interpretation Hnf4a’s role in the adult liver. In an earlier study (9), an increase in hepatocyte proliferation was observed following tamoxifen treatment of *Alb*-CreER^T2^; *Hnf4a^loxP/loxP^* mice, a result that was interpreted as indicating that Hnf4a has tumor suppressive (cell cycle inhibitor) activity. With the benefit of lineage tracing, by contrast, our experiments reveal reduced fitness of Hnf4a-deficient hepatocytes, resulting in their replacement by *Hnfa* wild-type cells (**Fig. 2**). Consequently, the increase in proliferation in Hnf4a-deficient livers may be due to the expansion of “escaper” cells rather than a cell-autonomous effect of Hnf4a.

A second advantage of the system is its ability to perform multi-gene knockout experiments. This can be achieved through one of two approaches: (i) simultaneous co-injection of multiple AAVs carrying individual gene-targeting sgRNAs or (ii) use of a polycistronic (PTG) AAV viral construct containing up to 4 independent sgRNAs. Using the co-injection strategy, we were able to demonstrate gene interactions between *Tfam*, *Polg,* and *Polg2* in mitochondrial maintenance (**Fig. S3**) and recapitulate features of *Lats1*/*Lats2* DKO mice (30) (**Fig. 5, Fig. S5, Fig. S6**). Although we observed similar findings with a polycistronic *Lats1*/*Lats2* PTG construct (**Fig. 5, Fig. S6**), the efficiency of multi-gene editing using the PTG strategy was lower than that achieved with the co-injection strategy, likely due to the requirement for simultaneous processing of 4 sgRNAs from a single template. Studies in plants and flies have reported that insertion of a tRNA between the PolIII promoter and the first sgRNA increased transcription efficiency (31), which was found to be due to the additive effect by the innate tRNA’s promoter activity (32). We attempted this approach without success (data not shown), an indication of species-specific differences in short RNA processing.

In summary, RIME represents a fast, cost-effective, and straightforward means of assessing conditional gene function *in vivo*. Moreover, the ability to perform multiplexed gene targeting provides an opportunity to assess questions of genetic epistasis and gene redundancy. As the spectrum of AAV capsids with tissue-specific tropism continues to evolve, this approach can be extended to other organs and organisms to address a wide range of biological questions without a need to create conditionally targeted ES cells or perform multiple rounds of breeding.

## Acknowledgements

We thank Jeffrey Posey for vectors and advice on viral preparation and members in the Stanger laboratory for comments and helpful suggestions. This work was supported by grants from the NIH (DK08355) and the Biesecker Pediatric Liver Foundation.

## Author contributions

Conceptualization: TK, BZS

Methodology: TK, KPS, MG, BZS

Experiments: TK, HC

Supervision: BZS

Discussion: ZA

Writing: TK, BZS

## Competing interests

None declared.

## Notes

### Competing Interest Statement

Dr. Grompe consults for, received grants from, and owns stock in Ambys and LogicBio. He consults for and owns stock in Yecuris.

## References

1. Wang T, Wei JJ, Sabatini DM, Lander ES. Genetic screens in human cells using the CRISPR-Cas9 system. Science. 2014;343:80–4.

2. Shalem O, Sanjana NE, Hartenian E, Shi X, Scott DA, Mikkelson T, et al. Genome-scale CRISPR-Cas9 knockout screening in human cells. Science. 2014;343:84–87.

3. Chen S, Sanjana NE, Zheng K, Shalem O, Lee K, Shi X, et al. Genome-wide CRISPR screen in a mouse model of tumor growth and metastasis. Cell. 2015;160:1246–1260.

4. Swiech L, Heidenreich M, Banerjee A, Habib N, Li Y, Trombetta J, et al. In vivo interrogation of gene function in the mammalian brain using CRISPR-Cas9. Nat. Biotechnol. 2015;33:102–106.

5. Platt RJ, Chen S, Zhou Y, Yim MJ, Swiech L, Kempton HR, et al. CRISPR-Cas9 knockin mice for genome editing and cancer modeling. Cell. 2014;159:440–455.

6. Soto F, Zhao L, Kerschensteiner D. Synapse maintenance and restoration in the retina by NGL2. Elife. 2018;7:1–15.

7. Laidlaw BJ, Duan L, Xu Y, Vazquez SE, Cyster JG. The transcription factor Hhex cooperates with the corepressor Tle3 to promote memory B cell development. Nat. Immunol. 2020;21:1082–1093.

8. Wei Y, Tian C, Zhao Y, Liu X, Liu F, Li S, et al. MRG15 orchestrates rhythmic epigenomic remodelling and controls hepatic lipid metabolism. Nat. Metab. 2020;2:447–460.

9. Bonzo JA, Ferry CH, Matsubara T, Kim JH, Gonzalez FJ. Suppression of hepatocyte proliferation by hepatocyte nuclear factor 4α in adult mice. J. Biol. Chem. 2012;287:7345–7356.

10. Yanger K, Zong Y, Maggs LR, Shapira SN, Maddipati R, Aiello NM, et al. Robust cellular reprogramming occurs spontaneously during liver regeneration. Genes Dev. 2013;27:719–24.

11. Tarlow BDD, Pelz C, Naugler WEE, Wakefield L, Wilson EMM, Milton J, et al. Bipotential Adult Liver Progenitors Are Derived from Chronically Injured Mature Hepatocytes. Cell Stem Cell. 2014;15:605–618.

12. Wang MJ, Chen F, Lau JTY, Hu YP. Hepatocyte polyploidization and its association with pathophysiological processes. Cell Death Dis. 2017;8:e2805.

13. Gentric G, Desdouets C. Polyploidization in liver tissue. Am. J. Pathol. 2014;184:322–331.

14. Wojcik EJ, Buckley RS, Richard J, Liu L, Huckaba TM, Kim S. Kinesin-5: Cross-bridging mechanism to targeted clinical therapy. Gene. 2013;531:133–149.

15. Chauvière M, Kress C, Kress M. Disruption of the mitotic kinesin Eg5 gene (Knsl1) results in early embryonic lethality. Biochem. Biophys. Res. Commun. 2008;372:513–519.

16. Chen F, Jimenez RJ, Sharma K, Luu HY, Hsu BY, Ravindranathan A, et al. Broad Distribution of Hepatocyte Proliferation in Liver Homeostasis and Regeneration. Cell Stem Cell. 2020;26:27–33.e4.

17. Katsuda T, Kawamata M, Hagiwara K, Takahashi R, Yamamoto Y, Camargo FD, et al. Conversion of Terminally Committed Hepatocytes to Culturable Bipotent Progenitor Cells with Regenerative Capacity. Cell Stem Cell. 2017;20:41–55.

18. Miyaoka Y, Ebato K, Kato H, Arakawa S, Shimizu S, Miyajima A. Hypertrophy and unconventional cell division of hepatocytes underlie liver regeneration. Curr. Biol. 2012;22:1166–75.

19. Gorman GS, Chinnery PF, DiMauro S, Hirano M, Koga Y, McFarland R, et al. Mitochondrial diseases. Nat. Rev. Dis. Prim. 2016;2:1–23.

20. Suomalainen A, Battersby BJ. Mitochondrial diseases: The contribution of organelle stress responses to pathology. Nat. Rev. Mol. Cell Biol. 2018;19:77–92.

21. Larsson NG, Wang J, Wilhelmsson H, Oldfors A, Rustin P, Lewandoski M, et al. Mitochondrial transcription factor A is necessary for mtDNA maintenance and embryogenesis in mice. Nat. Genet. 1998;18:231–236.

22. Hance N, Ekstrand MI, Trifunovic A. Mitochondrial DNA polymerase gamma is essential for mammalian embryogenesis. Hum. Mol. Genet. 2005;14:1775–1783.

23. Humble MM, Young MJ, Foley JF, Pandiri AR, Travlos GS, Copeland WC. Polg2 is essential for mammalian embryogenesis and is required for mtDNA maintenance. Hum. Mol. Genet. 2013;22:1017–1025.

24. Ekstrand MI, Falkenberg M, Rantanen A, Park CB, Gaspari M, Hultenby K, et al. Mitochondrial transcription factor A regulates mtDNA copy number in mammals. Hum. Mol. Genet. 2004;13:935–944.

25. El-Hattab AW, Craigen WJ, Scaglia F. Mitochondrial DNA maintenance defects. Biochim. Biophys. Acta - Mol. Basis Dis. 2017;1863:1539–1555.

26. Do Y, Matsuda S, Inatomi T, Nakada K, Yasukawa T, Kang D. The accessory subunit of human DNA polymerase γ is required for mitochondrial DNA maintenance and is able to stabilize the catalytic subunit. Mitochondrion. 2020;53:133–139.

27. Kim TY, Wangs D, Kim AK, Laus E, Lin AJ, Liem DA, et al. Metabolic labeling reveals proteome dynamics of mouse mitochondria. Mol. Cell. Proteomics. 2012;11:1586–1594.

28. Craven L, Alston CL, Taylor RW, Turnbull DM. Recent Advances in Mitochondrial Disease. Annu. Rev. Genomics Hum. Genet. 2017;18:257–275.

29. Chen Q, Zhang N, Xie R, Wang W, Cai J, Choi KS, et al. Homeostatic control of Hippo signaling activity revealed by an endogenous activating mutation in YAP. Genes Dev. 2015;29:1285–1297.

30. Yimlamai D, Christodoulou C, Galli GG, Yanger K, Pepe-Mooney B, Gurung B, et al. Hippo pathway activity influences liver cell fate. Cell. 2014;157:1324–38.

31. Xie K, Minkenberg B, Yang Y, S-s SIT. Boosting CRISPR/Cas9 multiplex editing capability with the endogenous tRNA-processing system. Proc. Natl. Acad. Sci. U. S. A. 2015;112:3570–3575.

32. Knapp DJHF, Michaels YS, Jamilly M, Ferry QRV, Barbosa H, Milne TA, et al. Decoupling tRNA promoter and processing activities enables specific Pol-II Cas9 guide RNA expression. Nat. Commun. 2019;10.

