## Supplemental Figures and Tables for "Rapid *in vivo* multiplexed editing (RIME) of the adult mouse liver"

**Supplementary Figure Legends**

**Figure S1. Characterization of sg*Hnf4a*-injected liver.**

(A) Location of each of the three sgRNAs targeting *Hnf4a*.

(B) qRT-PCR of *Hnf4a* with primers designed at the proximity of cut sites of each sgRNA.

(C) Gene expression analysis of hepatic marker genes by qRT-PCR ($n = 4$ at each time point). The data are normalized with *Gapdh*.

Statistical differences were calculated using multiple t-test. *$p < 0.05$, **$p < 0.01$, ***$p < 0.001$, ****$p < 0.0001$

### Figure S1

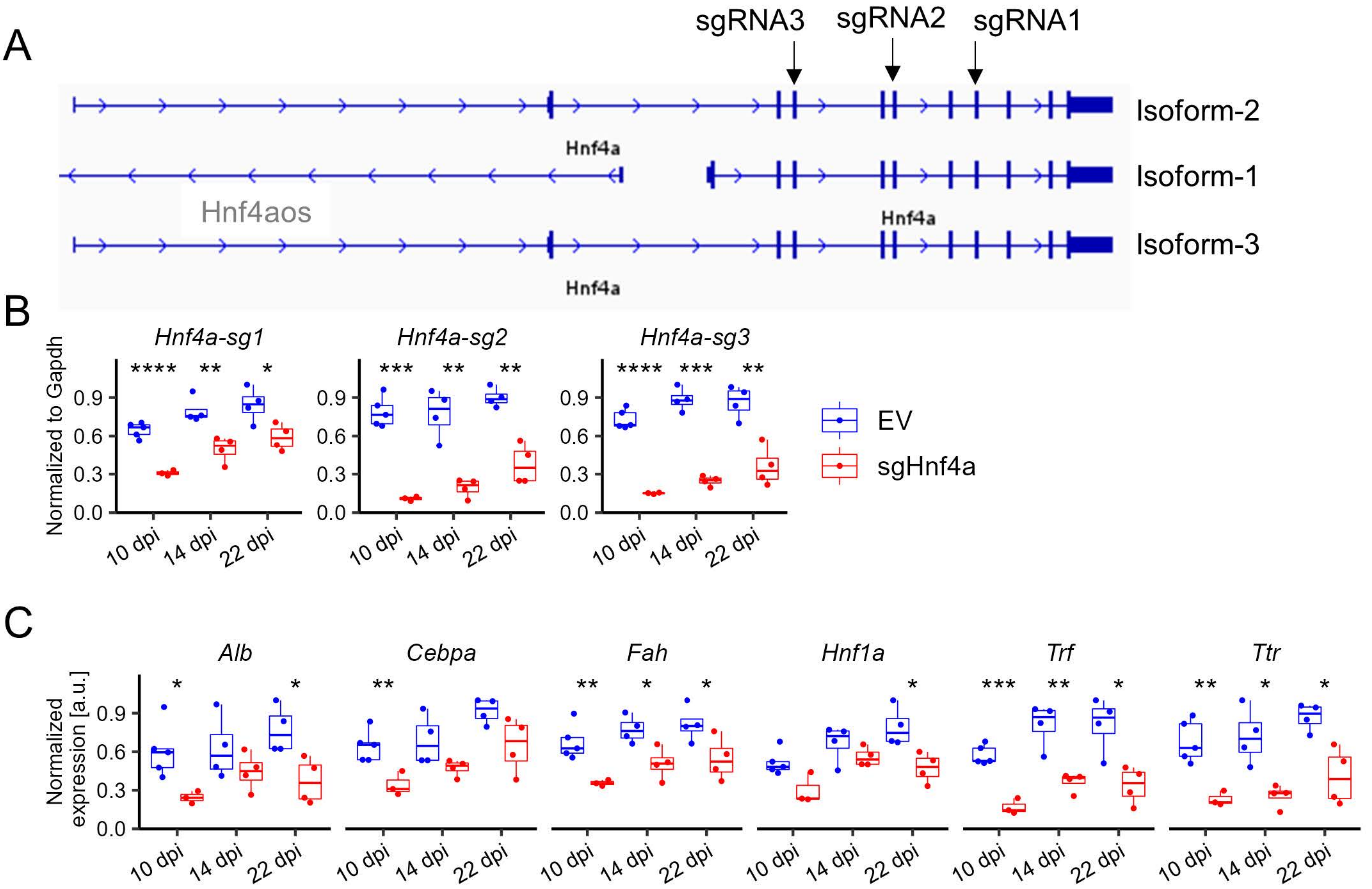
A
sgRNA3
sgRNA2
sgRNA1
Isoform-2
Hnf4a
Isoform-1
Hnf4aos
Hnf4a
Isoform-3
Hnf4a
B
Hnf4a-sg1
Hnf4a-sg2
Hnf4a-sg3
Normalized to Gapdh
0.0
0.3
0.6
0.9
10 dpi
14 dpi
22 dpi
**** ** *
*** ** **
**** *** **
EV
sgHnf4a
C
Alb
Cebpa
Fah
Hnf1a
Trf
Ttr
Normalized expression [a.u.]
* *
**
** * *
*
*** ** *
** * *

**Figure S2. Characterization of *Kif11*-LKO mice.**

(A) Representative flow cytometry results for nuclear DNA content by DAPI staining of isolated nuclei harvested from mice injected with EV or *sgKif11*.

(B) Quantification of the panel (A) (n = 3-5 for each data point). Stacked bar plots on the far right indicate the distribution of the nuclear DNA contents.

(C) Liver to body weight ratios were measured for *sgKif11*-injected mice harvested at 13 wpi. Age-matched mice without AAV injection were used as the control.

(D) GFP+ hepatocyte rates were estimated by flow cytometry at the indicate time points (n = 3-5 except for *sgKif11*-injected mice at 24 wpi, for which 2 mice were used).

### Figure S2

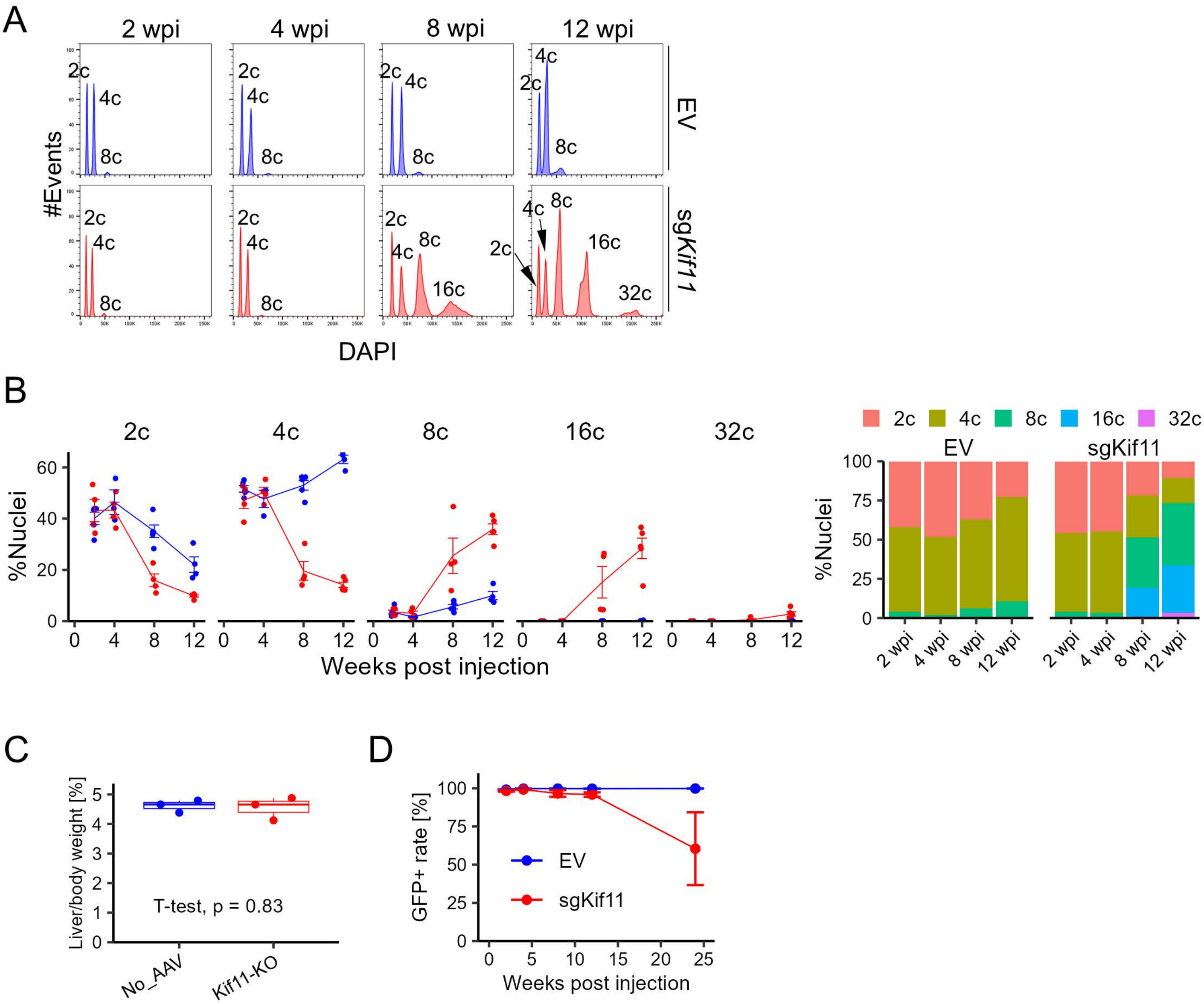
A
2 wpi
4 wpi
8 wpi
12 wpi
EV
sgKif11
#Events
2c
4c
8c
16c
32c
DAPI
B
2c
4c
8c
16c
32c
%Nuclei
Weeks post injection
EV
sgKif11
2 wpi
4 wpi
8 wpi
12 wpi
C
Liver/body weight [%]
T-test, p = 0.83
No_AAV
Kif11-KO
D
GFP+ rate [%]
EV
sgKif11
Weeks post injection

**Figure S3. Characterization of *Tfam*-, *Polg*- and *Polg2*-LKO mice.**

(A) Representative chromatograph of Sanger sequenced *Polg2* PCR amplicons from hepatocytes injected with EV, sg*Tfam*, sg*Polg* or sg*Polg2* to confirm *Polg2* liver knockout.

(B) Western blots of Tfam and Polg at the designated time points from livers injected with EV, sg*Tfam*, sg*Polg* or sg*Polg2*.

(C) qRT-PCR of *Tfam*, *Polg* and *Polg2*. Primers were designed randomly for *Tfam* and *Polg*, while each of the 3 *Polg2*-sgRNAs was used as one of the primers for *Polg2* quantification.

(D) Time course of the emergence of GFP+ hepatocytes as assessed by flow cytometry at 2, 4, 6, 8, 10 and 13 dpi (n = 2-7).

(E) Blood biochemistry at 10 and 11 wpi (n = 4) for EV and sg*Polg2*-injected mice. The data for 10 wpi are the same as those presented in **Fig. 4F**. The units on the y-axes are U/L for ALP, ALT and AST, µM for Bile Acids, and mg/dL for Cholesterol and Total bilirubin.

(F) Body weights at 13 wpi of mice injected with EV, sg*Tfam*, sg*Polg* sg*Polg2*, sg*Polg*/sg*Polg2* and sg*Tfam*/sg*Polg*/sg*Polg2*. sg*Tfam*/sg*Polg*/sg*Polg2* died at 9-10 wpi (3 out of 3 mice).

### Figure S3

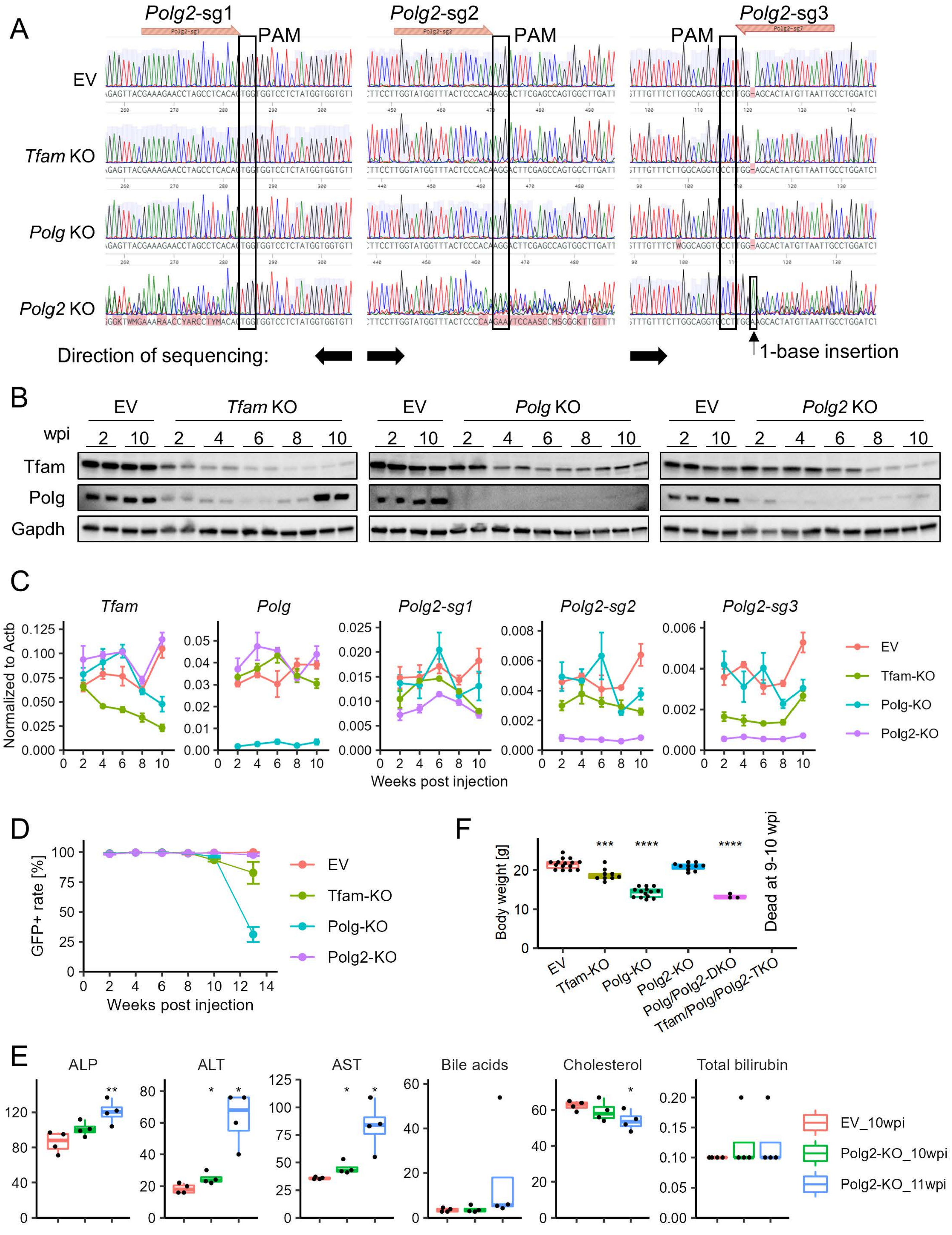
A
Polg2-sg1
Polg2-sg2
Polg2-sg3
PAM
EV
Tfam KO
Polg KO
Polg2 KO
Direction of sequencing:
1-base insertion
B
EV
Tfam KO
Polg KO
Polg2 KO
wpi
Tfam
Polg
Gapdh
C
Tfam
Polg
Polg2-sg1
Polg2-sg2
Polg2-sg3
Normalized to Actb
Weeks post injection
EV
Tfam-KO
Polg-KO
Polg2-KO
D
GFP+ rate [%]
Weeks post injection
F
Body weight [g]
Dead at 9-10 wpi
EV
Tfam-KO
Polg-KO
Polg2-KO
Polg/Polg2-DKO
Tfam/Polg/Polg2-TKO
E
ALP
ALT
AST
Bile acids
Cholesterol
Total bilirubin
EV_10wpi
Polg2-KO_10wpi
Polg2-KO_11wpi

**Figure S4. Characterization of *EGFP*/*Hnf4a* DKO by co-injection or PTG targeting.**

(A) Schematic representation of the strategy of DKO of *EGFP*/*Hnf4a* by co-injection of 2 AAVs (left) and by injection of a polycistronic AAV-PTG-*EGFP*/*Hnf4a* (right).

(B) Representative macroscopic images of *EGFP*/*Hnf4a* DKO at the designated time points.

(C) Schematic of the PTG-*EGFP*/*Hnf4a* vector

(D) Western blots of EGFP and Hnf4a at the designated time points.

(E) Flow cytometric quantification of GFP+ rates and GFP mean fluorescent intensity (MFI) of GFP+ hepatocytes at 28 dpi.

Statistical differences were calculated using multiple t-test. *$p < 0.05$, ***$p < 0.001$.

### Figure S4

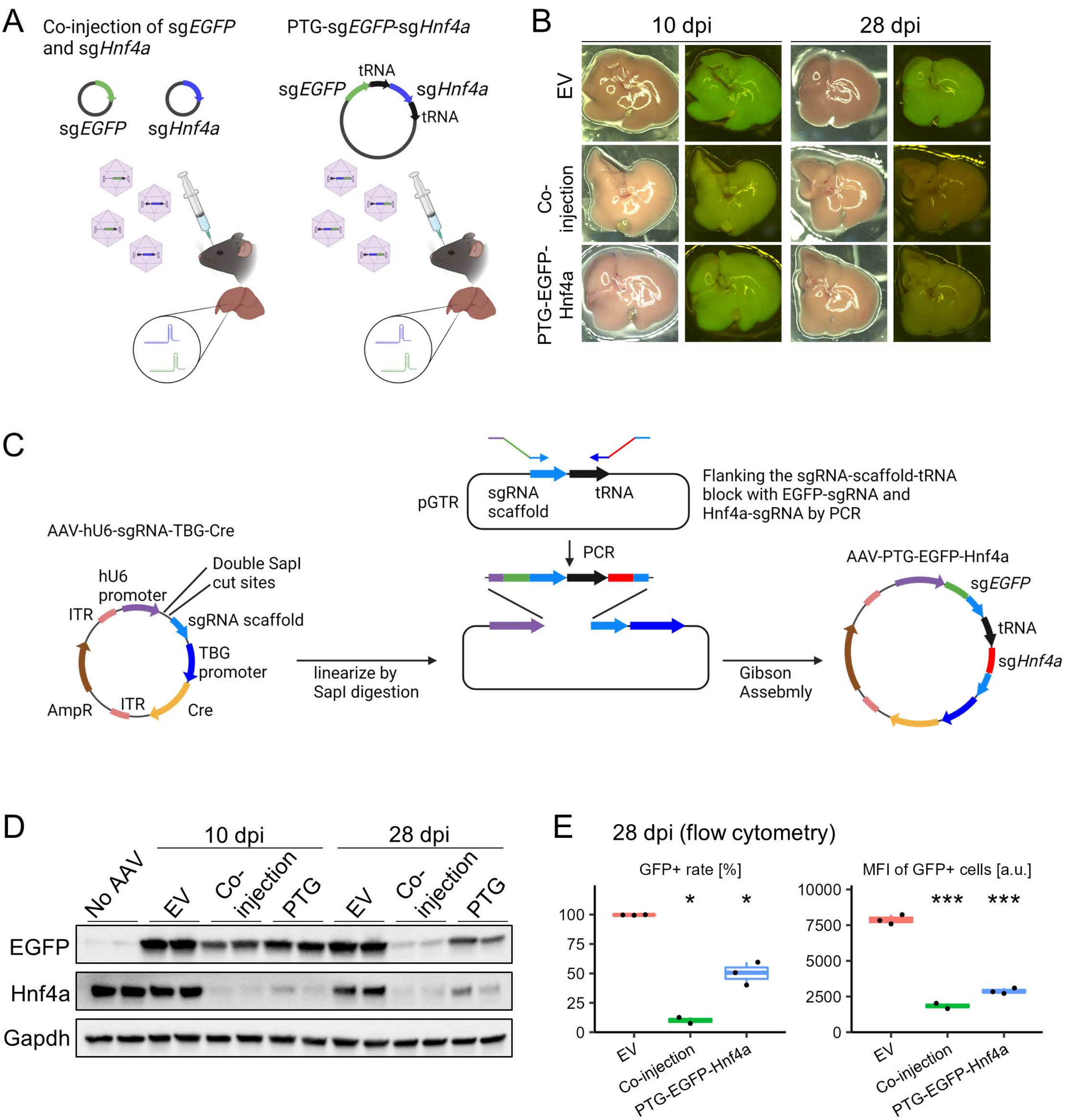
A
Co-injection of sgEGFP and sgHnf4a
sgEGFP
sgHnf4a
PTG-sgEGFP-sgHnf4a
tRNA
sgEGFP
sgHnf4a
tRNA
B
10 dpi
28 dpi
EV
Co-injection
PTG-EGFP-Hnf4a
C
AAV-hU6-sgRNA-TBG-Cre
hU6 promoter
Double SapI cut sites
ITR
sgRNA scaffold
TBG promoter
AmpR
ITR
Cre
linearize by SapI digestion
pGTR
sgRNA scaffold
tRNA
Flanking the sgRNA-scaffold-tRNA block with EGFP-sgRNA and Hnf4a-sgRNA by PCR
PCR
Gibson Assebmly
AAV-PTG-EGFP-Hnf4a
sgEGFP
tRNA
sgHnf4a
D
10 dpi
28 dpi
No AAV
EV
Co-injection
PTG
EV
Co-injection
PTG
EGFP
Hnf4a
Gapdh
E
28 dpi (flow cytometry)
GFP+ rate [%]
100
75
50
25
0
*
*
EV
Co-injection
PTG-EGFP-Hnf4a
MFI of GFP+ cells [a.u.]
10000
7500
5000
2500
0
***
***
EV
Co-injection
PTG-EGFP-Hnf4a

**Figure S5. Schematic representation of the cloning strategies for PTG.**

(A) Representative macroscopic images of the livers injected with EV, AAV-*Lats1-sg1/2*, AAV-*Lats2-sg1/2* and the liver co-injected with AAV-*Lats1-sg1/2* and AAV-*Lats2-sg1/2*.

(B) Schematic of the PTG-*Lats1*/*2* vector

(C) Chromatograph of bulk Sanger sequencing for each of the cut sites targeted by 2 *Lats2* sgRNAs.

### Figure S5

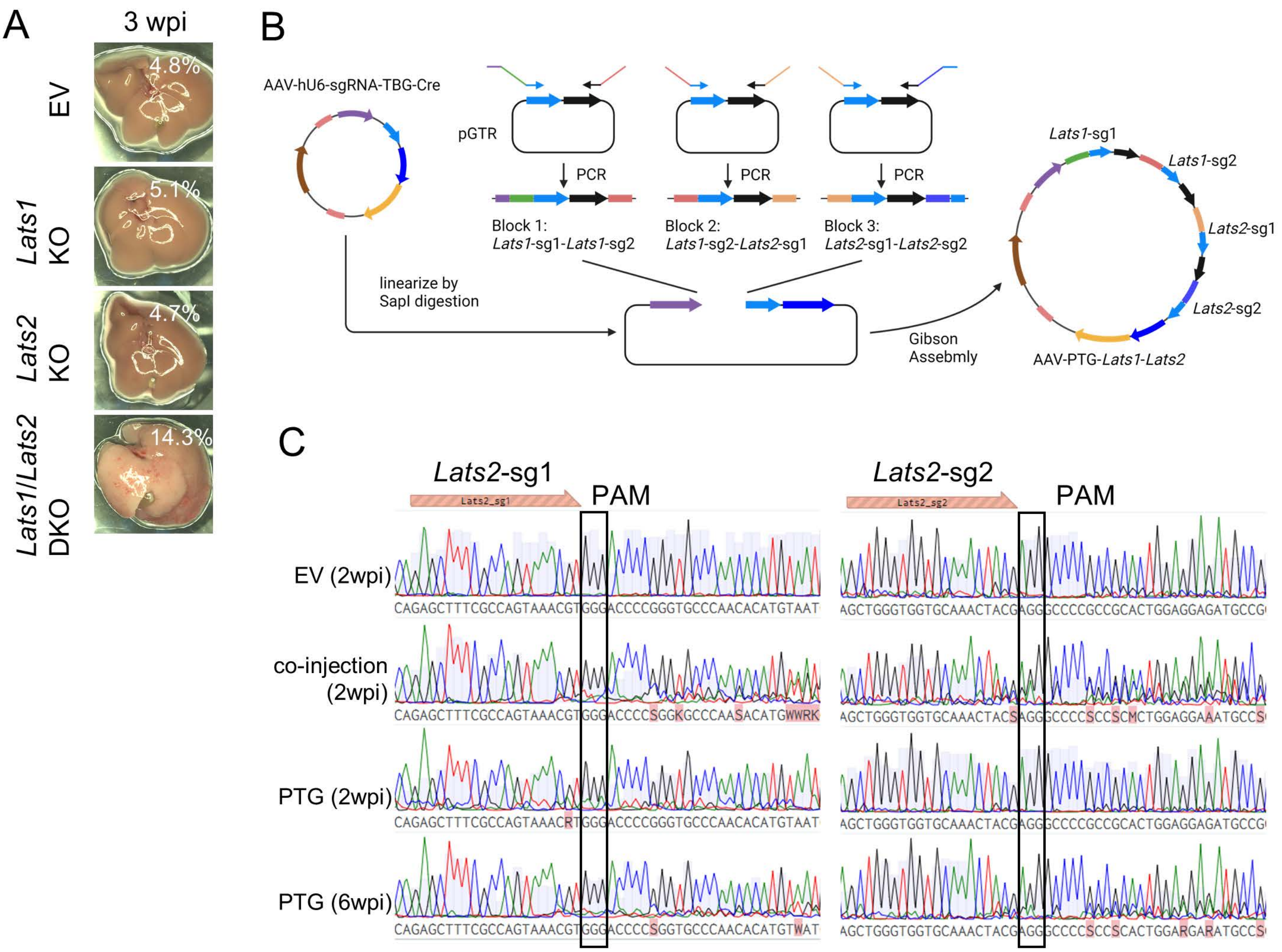
A
3 wpi
EV
4.8%
Lats1 KO
5.1%
Lats2 KO
4.7%
Lats1/Lats2 DKO
14.3%
B
AAV-hU6-sgRNA-TBG-Cre
pGTR
PCR
PCR
PCR
Block 1:
Lats1-sg1-Lats1-sg2
Block 2:
Lats1-sg2-Lats2-sg1
Block 3:
Lats2-sg1-Lats2-sg2
linearize by
SapI digestion
Gibson
Assebmly
Lats1-sg1
Lats1-sg2
Lats2-sg1
Lats2-sg2
AAV-PTG-Lats1-Lats2
C
Lats2-sg1
PAM
Lats2-sg2
PAM
EV (2wpi)
co-injection
(2wpi)
PTG (2wpi)
PTG (6wpi)

**Figure S6. Characterization of *Lats1*/*Lats2* DKO by co-injection and PTG.**

HE staining (top row) and immunofluorescence images for Sox9 and GFP (middle row) and Krt19 and GFP (bottom row). For immunofluorescence, nuclei were counterstained with DAPI.

### Figure S6

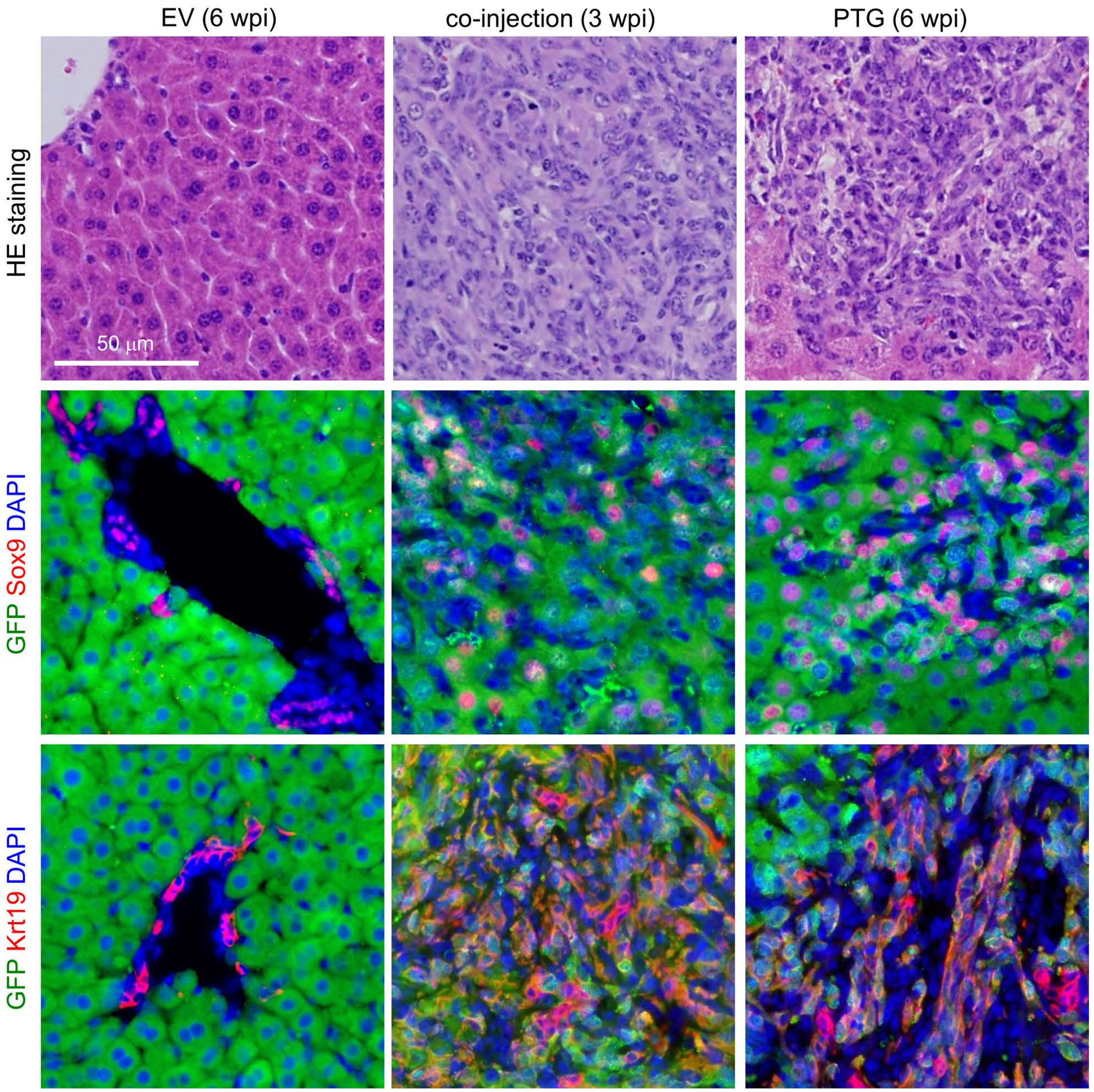

EV (6 wpi)
co-injection (3 wpi)
PTG (6 wpi)
HE staining
50 μm
GFP Sox9 DAPI
GFP Krt19 DAPI

#### Supplementary Tables

**Table S1. Oligo DNAs for sgRNA cloning.**

| sgRNA | Sense | Anti-sense |
|---|---|---|
| *EGFP* | ACCGGAAGTTCGAGGGCGACACCC | AACGGGTGTCGCCCTCGAACTTCC |
| *Hnf4a-sg1* | ACCGGCAATGACTACATCGTCCCT | AACAGGGACGATGTAGTCATTGCC |
| *Hnf4a-sg2* | ACCGGGGACCGGATCAGCACGCGG | AACCCGCGTGCTGATCCGGTCCCC |
| *Hnf4a-sg3* | ACCGCAGGTTTAGCCGACAATGTG | AACCACATTGTCGGCTAAACCTGC |
| *Kif11-sg1* | ACCGAGAAATCTGAGAAACACGAG | AACCTCGTGTTTCTCAGATTTCTC |
| *Kif11-sg2* | ACCGAACCACACTAGGGGCTACGA | AACTCGTAGCCCCTAGTGTGGTTC |
| *Kif11-sg3* | ACCGGTCACATTCCACTACTGAGT | AACACTCAGTAGTGGAATGTGACC |
| *Tfam-sg1* | ACCGTAAATGTTATATGCTGAACG | AACCGTTCAGCATATAACATTTAC |
| *Tfam-sg2* | ACCGGGAGCGTGCTAAAAGCACTG | AACCAGTGCTTTTAGCACGCTCCC |
| *Tfam-sg3* | ACCGCGGCCTACCTGATAGACGAG | AACCTCGTCTATCAGGTAGGCCGC |
| *Polg-sg1* | ACCGCCATACCTGGTCACAACCCG | AACCGGGTTGTGACCAGGTATGGC |
| *Polg-sg2* | ACCGTGCTCATAAACGTATCAGGT | AACACCTGATACGTTTATGAGCAC |
| *Polg-sg3* | ACCGCCGTTGCCATGGTGATACGT | AACACGTATCACCATGGCAACGGC |
| *Polg2-sg1* | ACCGGAAAGAACCTAGCCTCACAG | AACCTGTGAGGCTAGGTTCTTTCC |
| *Polg2-sg2* | ACCGTGGTATGGTTTACTCCCACA | AACTGTGGGAGTAAACCATACCAC |
| *Polg2-sg3* | ACCGCCTTAATAGGGGCTAAACAA | AACTTGTTTAGCCCCTATTAAGGC |
| *Lats1-sg1* | ACCGGGCAACCTAACATACCAGTG | AACCACTGGTATGTTAGGTTGCCC |
| *Lats1-sg2* | ACCGGCAGACATCTGCTCTCGACG | AACCGTCGAGAGCAGATGTCTGCC |
| *Lats2-sg1* | ACCGGAGCTTTCGCCAGTAAACGT | AACACGTTTACTGGCGAAAGCTCC |
| *Lats2-sg2* | ACCGGCTGGGTGGTGCAAACTACG | AACCGTAGTTTGCACCACCCAGCC |

**Table S2. PCR primers used for PTG cloning by Gibson assembly.**

| Construct | Block Fw/Rv | Primer sequence |
|---|---|---|
| PTG-*EGFP*-sg1-*Hnf4a*-sg2 | Block-1 Fw | TGGAAAGGACGAAACACCGGAAGTTCGAGGGCGACACCCG TTTTAGAGCTAGAAATAGCAAGTTAAAATAAGGC |
| | Block-1 Rv | GCTATTTCTAGCTCTAAAACCCGCGTGCTGATCCGGTCCCC TGCACCAGCCGGGAAT |
| PTG-*Lats1*-sg1-*Lats1*-sg2-*Lats2*-sg1-*Lats2-sg2* | Block-1 Fw | TGGAAAGGACGAAACACCGGGCAACCTAACATACCAGTGG TTTTAGAGCTAGAAATAGCAAGTTAAAATAAGGC |
| | Block-1 Rv | CGTCGAGAGCAGATGTCTGCCTGCACCAGCCGGGAAT |
| | Block-2 Fw | GGCAGACATCTGCTCTCGACGGTTTTAGAGCTAGAAATAGC AAGTTAAAATAAGGC |
| | Block-2 Rv | ACGTTTACTGGCGAAAGCTCCTGCACCAGCCGGGAAT |
| | Block-3 Fw | GGAGCTTTCGCCAGTAAACGTGTTTTAGAGCTAGAAATAGC AAGTTAAAATAAGGC |
| | Block-3 Rv | GCTATTTCTAGCTCTAAAACCGTAGTTTGCACCACCCAGCC TGCACCAGCCGGGAAT |

**Table S3. Primers used for qPCR.**

| Target | Forward | Reverse |
|---|---|---|
| AAV titration | GGAACCCCTAGTGATGGAGTT | CGGCCTCAGTGAGCGA |
| *Hnf4a*-sg1 | CCTCGGCACTGTCCAGAG | CATTGTCATCAATCTGCAGCTC |
| *Hnf4a*-sg2 | CGGAGGTCAAGCTACGAGGACA | TGATCCCAGAGATGGGAGAGGTG |
| *Hnf4a*-sg3 | GTGTGGTAGACAAAGATAAGAGG | CATTTTGGACAGCTTCCTTCT |
| *Epcam* | TCTACAAGGAAGAAATCAGCAAAA | CCCTCCTCAGTTCAGCACTC |
| *Hes1* | ACCGGCAATGACTACATCGTCCCT | AACAGGGACGATGTAGTCATTGCC |
| *Hnf1b* | TCTCACCAGCATGTCTTCCA | AAAATGGGGTCCTTGTTGCT |
| *Sox4* | CCTCGCTCTCCTCGTCCT | TCGTCTTCGAACTCGTCGT |
| *Sox9* | GACTCCCCACATTCCTCCTC | CCCTCTCGCTTCAGATCAAC |
| *Spp1* | GCTTGGCTTATGGACTGAGG | CGCTCTTCATGTGAGAGGTG |
| *Alb* | GCTGAGACCTTCACCTTCCA | CTTGTGCTTCACCAGCTCAG |
| *Cebpa* | CTCCCAGAGGACCAATGAAA | AAGTCTTAGCCGGAGGAAGC |
| *Fah* | CGGCGATGAAGTCATCATAA | GAGCTTCAGGCTGGTGAAAG |
| *Hnf1a* | GTCGAACATCCAGCACCTG | CCGTTGGAGTCGGAACTCT |
| *Trf* | TAGGAGCGGAGTACATGCAA | GAGCATCTGTCTCCACCACA |
| *Ttr* | TGGACACCAAATCGTACTGG | CAGAGTCGTTGGCTGTGAAA |
| *Polg2*-sg1 | ACCGGAAAGAACCTAGCCTCACAG* | CTGAAGGCACTGTCCCTCG |
| *Polg2*-sg2 | ACCGTGGTATGGTTTACTCCCACA* | AATCAGCGCTGCTGAAGTTAG |
| *Polg2*-sg3 | CAAGACAGAGAGCCGAGTAAG | AACTTGTTTAGCCCCTATTAAGGC* |
| *Tfam* | GGTCGCATCCCCTCGTCTATCA | GCTTCTGGTAGCTCCCTCCACA |
| *Polg* | AACTGGGCTGCTTAGACGTA | ATAAGGTCCGTTGCCATGGT |
| *Hk2* | GCCAGCCTCTCCTGATTTTAGTGT | GGGAACACAAAAGACCTCTTCTGG |
| *mt-Nd1* | CTAGCAGAAACAAACCGGGC | CCGGCTGCGTATTCTACGTT |
| *mt-Rnr2* | GCCAGCCTCTCCTGATTTTAGTGT | GGGAACACAAAAGACCTCTTCTGG |

*Oligo DNAs for sgRNA assembly were used as one of the primers.

**Table S4. Antibodies used for Western blotting.**

| Antibody | Host animal | Catalog # | Dilution | Manufacturer | Detection reagent |
|---|---|---|---|---|---|
| GFP | Goat | ab6673 | 1:2000 | Abcam | ECL |
| Gapdh | Rabbit | 2118 | 1:5000 | Cell Signaling | ECL Plus |
| Hnf4a | Rabbit | 3113 | 1:1000 | Cell Signaling | ECL Plus |
| Kif11 | Mouse | 627801 | 1:1000 | BioLegend | ECL |
| Tfam | Rabbit | ab131607 | 1:1000 | Abcam | ECL Plus |
| Polg | Rabbit | ab128899 | 1:1000 | Abcam | ECL Plus |
| Lats1 | Rabbit | 3477 | 1:5000 | Cell Signaling | West Femto |
| Yap/Taz | Rabbit | 8418 | 1:1000 | Cell Signaling | ECL Plus |
| pYap | Rabbit | 4911 | 1:1000 | Cell Signaling | ECL Plus |
| pTaz | Rabbit | 59971 | 1:1000 | Cell Signaling | ECL Plus |

**Table S5. Antibodies used for immunofluorescence.**

| Antibody | Host animal | Catalog # | Dilution | Manufacturer |
|---|---|---|---|---|
| GFP | Goat | ab6673 | 1:500 | Abcam |
| Hnf4a | Rabbit | 3113 | 1:200 | Cell Signaling |
| Yap/Taz | Rabbit | 8418 | 1:200 | Cell Signaling |
| Ki67 | Rat | 14-5698-82 | 1:200 | eBioscience |
| Sox9 | Rabbit | ab5535 | 1:200 | Millipore |
| Krt19 | Rabbit | N/A | 1:1000 | In-house |

**Table S6. Primers used for qPCR.**

| Target | Forward | Reverse |
|---|---|---|
| AAV titration | GGAACCCCTAGTGATGGAGTT | CGGCCTCAGTGAGCGA |
| *Hnf4a*-sg1 | CCTCGGCACTGTCCAGAG | CATTGTCATCAATCTGCAGCTC |
| *Hnf4a*-sg2 | CGGAGGTCAAGCTACGAGGACA | TGATCCCAGAGATGGGAGAGGTG |
| *Hnf4a*-sg3 | GTGTGGTAGACAAAGATAAGAGG | CATTTTGGACAGCTTCCTTCT |
| *Epcam* | TCTACAAGGAAGAAATCAGCAAAA | CCCTCCTCAGTTCAGCACTC |
| *Hes1* | ACCGGCAATGACTACATCGTCCCT | AACAGGGACGATGTAGTCATTGCC |
| *Hnf1b* | TCTCACCAGCATGTCTTCCA | AAAATGGGGTCCTTGTTGCT |
| *Sox4* | CCTCGCTCTCCTCGTCCT | TCGTCTTCGAACTCGTCGT |
| *Sox9* | GACTCCCCACATTCCTCCTC | CCCTCTCGCTTCAGATCAAC |
| *Spp1* | GCTTGGCTTATGGACTGAGG | CGCTCTTCATGTGAGAGGTG |
| *Alb* | GCTGAGACCTTCACCTTCCA | CTTGTGCTTCACCAGCTCAG |
| *Cebpa* | CTCCCAGAGGACCAATGAAA | AAGTCTTAGCCGGAGGAAGC |
| *Fah* | CGGCGATGAAGTCATCATAA | GAGCTTCAGGCTGGTGAAAG |
| *Hnf1a* | GTCGAACATCCAGCACCTG | CCGTTGGAGTCGGAACTCT |
| *Trf* | TAGGAGCGGAGTACATGCAA | GAGCATCTGTCTCCACCACA |
| *Ttr* | TGGACACCAAATCGTACTGG | CAGAGTCGTTGGCTGTGAAA |
| *Polg2*-sg1 | ACCGGAAAGAACCTAGCCTCACAG* | CTGAAGGCACTGTCCCTCG |
| *Polg2*-sg2 | ACCGTGGTATGGTTTACTCCCACA* | AATCAGCGCTGCTGAAGTTAG |
| *Polg2*-sg3 | CAAGACAGAGAGCCGAGTAAG | AACTTGTTTAGCCCCTATTAAGGC* |
| *Tfam* | GGTCGCATCCCCTCGTCTATCA | GCTTCTGGTAGCTCCCTCCACA |
| *Polg* | AACTGGGCTGCTTAGACGTA | ATAAGGTCCGTTGCCATGGT |
| *Hk2* | GCCAGCCTCTCCTGATTTTAGTGT | GGGAACACAAAAGACCTCTTCTGG |
| *mt-Nd1* | CTAGCAGAAACAAACCGGGC | CCGGCTGCGTATTCTACGTT |
| *mt-Rnr2* | GCCAGCCTCTCCTGATTTTAGTGT | GGGAACACAAAAGACCTCTTCTGG |

*Oligo DNAs for sgRNA assembly were used as one of the primers.
